# Computational and structure-based analysis of the conserved 5′–3′ exoribonuclease (XRN) family in plant species suggests seven potentially functional amino acid motifs within the N-terminal catalytic core

**DOI:** 10.64898/2026.11.26.769122

**Authors:** Isabel Stark, Christian G. Schwarz, Elias Wippermann, Till Rudack, Aline Koch

## Abstract

Plant diseases and pests cause substantial crop losses worldwide, underscoring the need for safe and effective plant protection measures. One potential approach is the foliar spraying of mRNA to induce transient protein expression in plants, but RNA instability limits its use. Plant 5′–3′ exoribonucleases (XRNs) could degrade exogenous RNA after uptake, thereby contributing to an intracellular barrier, while viral exoribonuclease-resistant RNAs (xrRNAs) could protect exogenous RNA. However, the molecular mechanisms remain unclear. Here, XRNs from *Arabidopsis thaliana*, *Zea mays*, *Hordeum vulgare*, *Nicotiana tabacum*, and *Nicotiana benthamiana* were investigated using computational methods such as phylogenetic, sequence-based and structure-based analyses. Phylogenetic analysis of eleven plant XRN candidates, together with the *A. thaliana* XRNs, grouped into two clades designated XRN4-like and XRN2/3-like, and an outgroup comprising two monocot sequences. In twelve of fifteen predicted plant XRN structures in complex with a 5 nucleotide (nt) single-stranded RNA (ssRNA) and two magnesium ions, the active center reproduced the architecture described for homologous XRN crystal structures. Contact and sequence analyses identified conserved amino acids within seven conserved motifs, six of which contacted ssRNA or magnesium ions in the models. No signal peptides indicative of apoplastic secretion were detected. Furthermore, modeling of a ST9a xrRNA-tagged ssRNA in complex with NbXRN4-like 1 showed that the xrRNA motif occupies a structural volume comparable to that of the XRN model, suggesting steric hindrance to substrate entry. These findings provide testable structural hypotheses for XRN-RNA interactions in plants and support computational modeling to investigate RNA stability and to develop more stable RNA molecules for plant applications.

**Graphical abstract:** 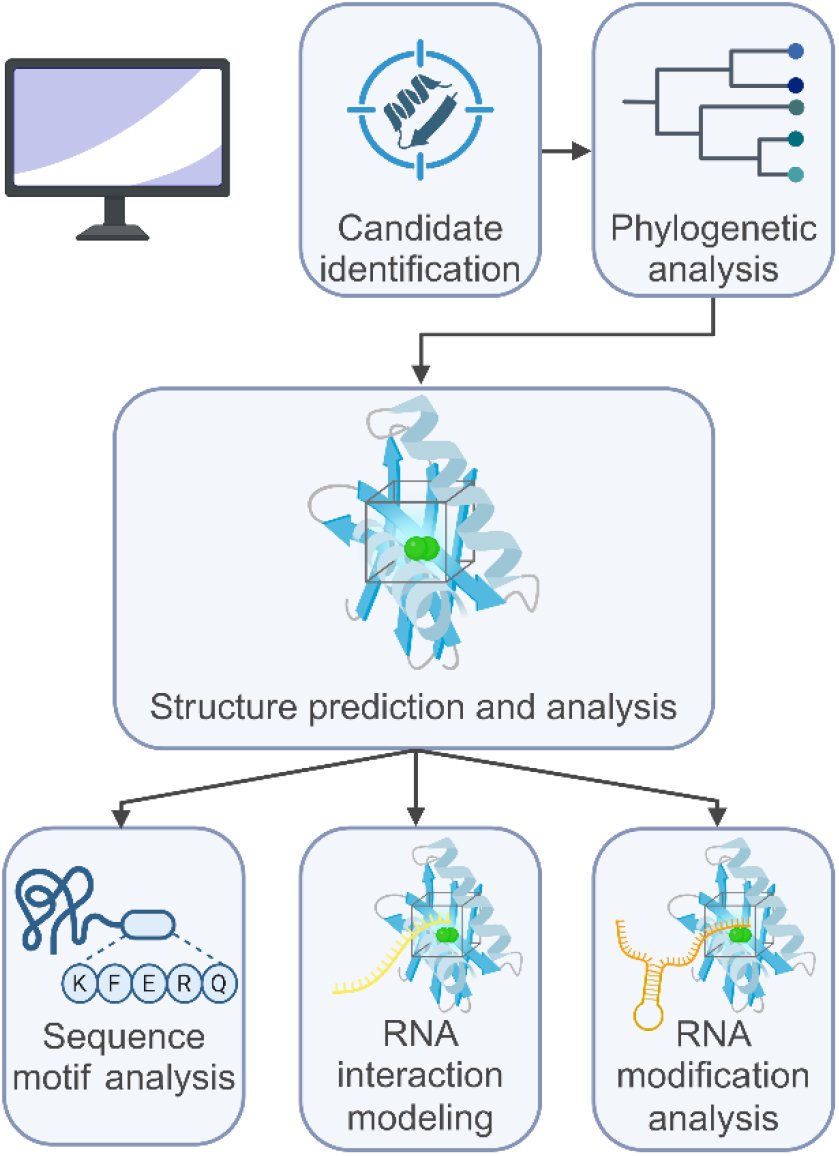

Created in BioRender. Schlemmer, T. (2026) https://BioRender.com/5gc8vfn

## Introduction

Every year, plant diseases and pests cause crop failures that result in financial losses of up to USD 220 billion [1]. This enormous damage to the global economy underscores the importance of plant protection measures. Currently, conventional pesticides are often used for this purpose, even though they and their by-products negatively impact human health and the environment [2,3]. Hence, society and politics are calling to reduce or completely eliminate the use of conventional pesticides [4,5]. Ensuring crop protection, however, requires fast-acting alternative strategies. One such approach is the spray application of RNA, with research in this field focusing on spray-induced gene silencing (SIGS) via RNA interference (RNAi). RNAi is a naturally occurring mechanism that selectively silences genes before their mRNA transcript is translated into protein, which is also known as post-transcriptional gene silencing (PTGS) [6]. It is primarily triggered by double-stranded RNA (dsRNA), which activates an enzymatic cascade that processes the dsRNA into small interfering RNAs (siRNAs), which then mediate the cleavage and ultimately the degradation of the target transcripts via the RNA-induced silencing complex (RISC) [7]. Although the effectiveness of dsRNA spray applications in combating fungal diseases has been demonstrated [8], some obstacles remain, such as the efficient cellular uptake of dsRNA molecules [9,10]. Another approach is the spray application of mRNA for transient expression of proteins inside plant cells, which could potentially enable the temporary restoration of gene function or the production of protective proteins [11]. For both concepts, RNA stability is a crucial aspect for high-performance spray applications.

To improve the stability of exogenous RNAs in plants, it is important to understand the molecular mechanisms of RNA-cleaving and RNA-degrading enzymes. These proteins are distributed throughout the cytoplasm, the nucleus, organelles, and the secretory pathway [12]. Although they perform multiple functions, they share a common purpose in maintaining the cellular RNA homeostasis [12]. One group of RNA-cleaving enzymes that deserves further attention in the context of RNA spray applications is the group of RNase three-like (RTL) proteins. They cleave dsRNA endonucleolytically, thereby suppressing siRNA production, which would reduce the overall efficiency of dsRNA spray applications [13]. RNA degradation might be a major limiting factor in RNA spray applications where ssRNAs such as mRNAs are applied. mRNA degradation is a well-documented pathway in eukaryotes that can be characterized by the direction of degradation mediated by its associated enzymes [14]. For example, the exosome complex degrades transcripts without poly(A)-tail in 3′–5′ direction [15]. On the other hand, 5′-monophosphorylated (5′-p) mRNAs are hydrolyzed in 5′–3′ direction by members of the highly conserved 5′–3′ exoribonuclease (XRN) family [14,16]. Therefore, understanding both molecular mechanisms in plants would hold great potential to improve RNA stability in RNA-based spray applications.

In plants, the XRNs from *A. thaliana* (Arabidopsis) have been investigated in more detail. Arabidopsis encodes three different XRNs, namely the nuclear AtXRN2 and AtXRN3, and the cytoplasmic AtXRN4, which is also known as ETHYLENE INSENSITIVE5 (EIN5) or ACC-INSENSITIVE1 (AIN1) [17]. AtXRN3 has two alternatively spliced isoforms and is crucial for viability and normal plant growth, as homozygous loss-of-function mutants show embryo lethality [18,19]. In contrast, loss of function of AtXRN2 results in no obvious developmental abnormalities [18], while loss of function of AtXRN4 causes only mild to no abnormalities under standard conditions [18,20]. However, the absence of the latter two can be measured at the molecular level. In absence of AtXRN2, specific defects occur during the cleavage of pre-rRNAs, which suggests that AtXRN2 participates in rRNA processing [21]. The absence of AtXRN4 leads to hormonal disorders in the plant, as this enzyme is responsible for the degradation of *EIN3-BINDING F-BOX PROTEIN 1* and *2* (*EBF1/2*) transcripts, which, once expressed, negatively regulate the ethylene signal transduction cascade [22–26]. Furthermore, AtXRN4 may act as a key regulator of the plant’s stress response, which is mediated by the stress hormone abscisic acid (ABA) [20,27]. Previous studies indicate that all Arabidopsis XRNs seem to act antagonistically in the PTGS pathway, with AtXRN4 showing the strongest effect [18,20,21,28–32]. For example, when AtXRN4 is absent, uncapped transcripts accumulate and serve as template for dsRNA production and, consequently, for the excessive production of siRNAs, which presumably leads to excessive silencing of genes such as *NITRATE REDUCTASE 1* and *2* (*NIA1/2*) [20]. These vital and diverse duties of Arabidopsis XRNs make the plant XRNs a promising research topic to increase efficacy of RNA spray applications.

Currently, important information about the specific architecture of plant XRNs is missing, since no crystal structures of these enzymes, either alone or in complex with a nucleotide substrate, are available. In other eukaryotic model systems, the architecture of members of the XRN family is well characterized. In particular, the enzyme-substrate interaction is resolved in the crystal structures of the cytoplasmic 5′–3′ exoribonuclease 1 (Xrn1p, called ScXRN1 hereafter) and the nuclear ribonucleic acid trafficking (Rat1p, called ScXRN2 hereafter) from *Saccharomyces cerevisiae*, as well as the ScXRN1 ortholog Pacman (called DmXRN1 hereafter) from *Drosophila melanogaster* [33–42]. All three crystal structures show the coordinated molecular interaction between the 5′-p substrate and the active center where three nucleotides are held in place by π-π-stacking between the aromatic side chains of the two conserved amino acids histidine (stacking His) and tryptophan (stacking Trp) [35,41,42]. Moreover, it is thought that two magnesium ions interact through direct or indirect coordination with the phosphate groups of the RNA backbone and with a group of six or seven amino acids consisting solely of aspartic acid (Asp), glutamic acid (Glu), and asparagine (Asn)[35,41]. In aqueous solution, one magnesium ion can directly interact with up to six ligands such as water molecules in an octahedral geometry [43]. Therefore, it is suggested that one of the magnesium ions provides a water molecule for hydrolysis [35]. Additionally, phylogenetic analysis suggests that ScXRN2 has the shortest phylogenetic distance to all Arabidopsis XRNs [21]. This knowledge can be utilized to model the structure of plant XRNs *in silico*, which would contribute to identifying ssRNA designs with increased resistance to degradation for spray applications. Here, possible ssRNA design motifs that protect the downstream area of ssRNAs are viral exoribonuclease-resistant RNA motifs (xrRNAs) [44–46]. For example, the plant-infecting beet western yellows virus ST9-associated (ST9a) RNA inhibits a member of the XRN family from *Kluyveromyces lactis in vitro* and modeling of this xrRNA’s crystal structure suggests the formation of a protective ring-structure around the 5′ end, which seems to sterically hinder exoribonuclease activity [44,47]. This viral motif could therefore be exploited to prevent degradation of exogenous ssRNAs. Overall, gaining a better understanding about how xrRNAs prevent RNA decay in plants holds opportunities for many biotechnological applications, such as heterologous or transient expression of proteins via ssRNA spray applications.

The aim of this study was the computational, structure- and sequence-based analysis of the XRN family in the plant species Arabidopsis, *Zea mays* (maize), *Hordeum vulgare* (barley), *Nicotiana tabacum,* and *Nicotiana benthamiana* (both commonly referred to as tobacco). In this study, phylogenetic analysis of eleven plant XRN candidates from maize, barley, *N. benthamiana*, and *N. tabacum*, together with *A. thaliana* XRNs, revealed two clades, designated XRN4-like and XRN2/3-like, and a separate outgroup comprising two monocot sequences. *In silico* structure prediction served as a tool to generate models of plant XRN protein structures in complex with a 5 nt long ssRNA and two magnesium ions. For twelve of the fifteen computational plant XRN models, the active center was predicted with two magnesium ions and the ssRNA in 5′ direction, which reproduced the active center of the homologous XRNs. Structure analysis of these twelve computational plant XRN models confirmed that four conserved amino acids within the active center interact with nt 1 or the magnesium ions. These amino acids were located within three of the seven conserved motifs specified by sequence motif analysis. Sequence motif analysis furthermore suggested that none of the identified plant XRNs are transported to the apoplast through signal peptides. To test the volume difference of the ST9a xrRNA motif [44] relative to the plant XRNs, a ST9a xrRNA-tagged ssRNA was modeled in complex with the NbXRN4-like 1 structure prediction model. It showed that the 3D xrRNA model occupies a structural volume much larger than the substrate entry site of the NbXRN4-like 1 model.

## Materials and methods

### Candidate screening

To identify XRN candidates in plants, basic local alignment search tool for proteins (BLASTP, version 2.16.0+ [48,49]) was used via the web server UniProt [50,51]. The full-length protein sequence of ScXRN2 (accession number Q02792) served as template, the E-value cutoff was set to 1 × 10⁻⁵, and the taxonomy was restricted to the following species: *A. thaliana* (species ID 3702), *N. tabacum* (species ID 4097), *H. vulgare* (species ID 4513), and *Z. mays* (species ID 4577). For Arabidopsis, previously published AtXRN2, the isoforms AtXRN3.1 and AtXRN3.2, and AtXRN4 were selected [17]. For all other species, protein sequences identified by BLASTP with an E-value below 1 × 10⁻¹⁰⁰ over the N-terminal region, and a predicted or assigned ribonuclease enzymatic function were included (Figure S 1). The identified sequences for *N. tabacum* were removed by UniProt, since they were not included in the UniProt reference genome (as of May 2026). Therefore, we included protein sequences from *N. benthamiana* and *N. tabacum* listed in UniProt as Similar Proteins with 90% sequence identity to the respective proteins from *N. tabacum*. The accession numbers of all protein sequences used are listed in Table S 1.

### Phylogenetic analysis

The evolutionary relationships among the selected protein sequences were analyzed in a phylogenetic context. For this purpose, a phylogenetic tree was generated via the web server NGPhylogeny [52,53]. Here, the protein sequences of ScXRN1, ScXRN2, DmXRN1, *Fadolivirus algeromassiliense* 5′–3′ exoribonuclease [54], and of the plant candidates were aligned with Clustal Omega [55]. The resulting multiple sequence alignment (MSA) was curated to be suitable for phylogenetic inference by Block Mapping and Gathering with Entropy (BMGE) [56]. Then, maximum likelihood-based inference of phylogenetic trees with Smart Model Selection (PhyML+SMS) was carried out, which ranks the substitution matrices depending on their similarity to the analyzed data and decides on the best one [57,58]. To test the robustness of the tree, the branch support was obtained by the phylogenetic bootstrap method with 1,000 bootstrap replicates. The phylogenetic tree was exported as Newick format [59] to the web server iTOL [60,61] to display the bootstraps as text.

### Plant XRN structure prediction and analysis

To predict protein structures in complex with a randomized 5 nt long ssRNA (5′-CCAUU-3′) and two magnesium ions, the following deep learning based structure prediction tools were used: both Chai-1 (version 0.6.2, [62]) and Boltz-2 (version 2.2.1, [63]) via the web server Tamarind Bio [64] as well as AlphaFold3 [65]. When structure prediction without modifications did not yield plausible results (selection criteria given below), the following optional settings were incorporated into the workflow of Chai-1 and Boltz-2 predictions: Due to its phylogenetic proximity, the crystal structure of ScXRN2 (PDB-ID 8K5P, [42]) was provided as template or contact restraints were added that specified the stacking His with nt 1 and the stacking Trp with nt 3 within a 5 Å distance. The generated multiple structure suggestions per XRN-RNA complex were analyzed in PyMOL [66] by aligning them to the crystal structures of ScXRN2, ScXRN1 (PDB-ID 6Q8Y, [41]), and DmXRN1 (PDB-ID 2Y35, [35]). To obtain plausible plant XRN models, out of the multiple structure suggestion models per XRN-RNA complex the best one was selected based on the following criteria mentioned in the literature [35,41,42]: Firstly, both RNA and magnesium ions had to be located within the same active center of the enzyme. Next, the orientation of the RNA was of crucial importance; therefore, the 5′ end had to be placed into the active center. Structure prediction models that assumed the formation of a loop in the middle of the RNA within the pocket were discarded. Lastly, the possibility of the stacking His and the stacking Trp interacting via π-π stacking with nt 1 and nt 3 respectively was examined. The selected structures were all modeled by Chai-1. In the following, only results from Chai-1 predictions of XRN-RNA complexes were used. The structure predictions were processed using MAXIMOBY (version 25.02 (12/25), [67]). Here, for each residue, the energies between Van-der-Waals clashes were resolved using the force field Amber84 [68] and conjugate gradient as optimizer. The structure was prepared by protonation using the p*K*_a_ (acid dissociation constant on a logarithmic scale) calculation based on Nielsen and Vriend [69]. Then, a residue level contact analysis based on geometric and chemical criteria was applied using kontakteUR and the resulting contact matrices were filtered for all contacts with magnesium ions or for XRN-RNA interactions [70]. The contacts were visualized using Gnuplot [71].

### Sequence motif analysis

Important sequence motifs were determined by scanning all sequences with Gapped Local Alignment of Motifs (GLAM2, version 1056 [72,73]). GLAM2 parameters were as follows: the search begins by aligning two initial columns and then continues up to a maximum of 20. For quality control, selected sequences were compared by MSA using Clustal Omega [74,75]. Additionally, signal peptide screening was performed to assess the potential secretion of plant XRNs into the apoplast. The protein sequences of all eukaryotic XRNs were submitted to SignalP-6.0 [76,77], with ScXRN1, ScXRN2 and DmXRN1 included for comparison.

### xrRNA-XRN complex modeling

To obtain a first computational model hypothesis of xrRNA in complex with a plant XRN in preparation for molecular dynamics simulation, a ST9a xrRNA-tagged ssRNA was modeled and fitted to the plant XRN model of NbXRN4-like 1. The tagged RNA sequence had a 60 nt long randomized leader sequence (5′-GGAAGAAUGA AACUGUCACC AUUUGUAUAA UAUUUCUAGU UAGGAUAGAG UCAUAACAAC-3′) upstream, which should enable the formation of the tertiary structure *in vitro*. The 3D structure of the tagged RNA was predicted via AlphaFold3 [65]. In accordance to the interacting regions of the crystal structure of ST9a xrRNA motif (PDB ID 9CFN, [44]), the structure prediction with the most interacting regions was selected, though the active conformation of the crystal structure was not reproduced. Therefore, the predicted ST9a xrRNA motif was substituted by the crystal structure as follows: In PyMOL, the stop site nucleotide of the ST9a xrRNA crystal structure was aligned to the corresponding nucleotide of the tagged RNA prediction. The stop site nucleotide of the crystal structure and the predicted ST9a xrRNA region were removed. The xrRNA model was then covalently bound to the ST9a xrRNA crystal structure and the energy of the bond was optimized in MOBY (version 25.02 (12/25), [67]). The resulting tagged-RNA structure prediction model, the first 42 nucleotides of the leader sequence were removed to obtain a region without hydrogen bonded nucleotides for active center fitting. The second nucleotides of the tagged-RNA structure model (G44) and of the ssRNA of NbXRN4-like 1 model (C2) were aligned.

### *In vitro* transcription (IVT)

To investigate the resistance of the ST9a xrRNA motif [44] to XRN in future RNA degradation assays, the ST9a xrRNA motif and the ST9a xrRNA motif carrying two point mutations (A12U/A13U) that are expected to impair tertiary structure formation were synthesized via IVT. Both versions of the xrRNA sequence had the same 60 nt long randomized leader sequence upstream, which should enable the formation of the tertiary structure. The DNA template for IVT was amplified by PCR and analyzed on a 1% (w/v) agarose gel. The PCR product was then purified using the NucleoSpin Gel and PCR Clean-up kit (MACHEREY-NAGEL, Düren, Germany; catalog number 740609.50) according to the manufacturer’s guidelines. IVT was carried out using the HiScribe T7 High Yield RNA Synthesis Kit (New England Biolabs, E2040) with 1 µg PCR product as template according to the manufacturer’s guidelines. After incubation overnight, 2 µL DNase I (Thermo Fisher Scientific, catalog number EN0525) were added, and the reaction was incubated at 37°C for 30 minutes to remove remaining DNA template. The generated RNA was purified using the Monarch Spin RNA Cleanup Kit (500 µg) (New England Biolabs, catalog number T2060) according to the manufacturer’s guidelines. RNA concentration was measured on a NanoDrop One^C^ Microvolume UV-Vis spectrophotometer (Thermo Fisher Scientific). To test the integrity of the *in vitro* transcribed RNA, 500 ng of RNA were analyzed on a 6% (w/v) urea polyacrylamide gel consisting of 6% (w/v) acrylamide in 50% (w/v) urea TBE buffer with 10% (w/v) ammonium persulfate (APS) and 0.1% (v/v) tetramethylethylenediamine (TEMED). Samples were prepared with 2X formamide loading dye and heated at 95°C for 10 min. GeneRuler Ultra Low Range DNA Ladder (Thermo Fisher Scientific, catalog number SM1211) was used as size standard. Electrophoresis was performed at 300 V for 30 min in 1X TBE buffer. After that, gels were submerged in a staining solution consisting of 1% (v/v) 20,000X ROTI GelStain (Carl ROTH, catalog number 3865) in 1X TBE buffer for 10 min. The fluorescent bands were visualized using the ChemiDoc Imaging System (Bio-Rad Laboratories, Hercules, CA, USA) set to ethidium bromide detection channel and rapid auto detect.

### PCR amplification

Sequences were amplified using Q5 High-Fidelity DNA Polymerase (New England Biolabs, catalog number M0491) in 50 µL reactions according to the manufacturer’s guidelines with the final extension time increased to 5 min. 300 ng of plasmid DNA (see Table S 2) served as template. Primers that were used are listed in Table S 3.

## Results

### Phylogenetic analysis reveals three distinct XRN clades in plants, two of which include the well-known XRNs from Arabidopsis

The XRN family is important for RNA homeostasis in all eukaryotes. Apart from Arabidopsis, very little is known about XRNs in other plant species. Therefore, it was necessary to identify possible plant XRN candidates and analyze their evolutionary relationships. For this purpose, BLAST-based candidate screening and phylogenetic analysis were conducted. Candidate screening revealed several XRN candidates for maize, barley and both Nicotiana species (Figure S 1), which were used for phylogenetic analysis. The plant XRNs clustered into two distinct clades designated XRN4-like clade after AtXRN4 and the XRN2/3-like clade after AtXRN2, AtXRN3.1, and AtXRN3.2, as well as a separate outgroup comprising two monocot sequences designated ZmXRN-like 1 and HvXRN-like 1 (Figure 1). The XRN4-like clade and the XRN2/3-like clade were classified as descending from a common ancestor. Within these two clades, the AtXRNs were grouped together with the other dicot XRNs from *N. benthamiana* and *N. tabacum*, resulting in a monocot subclade and a dicot subclade within each clade. Furthermore, the phylogenetic tree showed that all plant XRNs shared a common ancestor and were most closely related to ScXRN2. Moreover, ScXRN2 shared a more recent common ancestor with all the plant XRNs than ScXRN1, DmXRN1, and FaXRN-like. Branch support was heterogeneous: all branches, except for the branch containing AtXRN2 and AtXRN3.1/3.2, were supported by bootstrap support above 60%, with eleven of the sixteen branches supported by values above 90%. These findings indicate that most of the identified XRN candidates were closely related either to AtXRN4 or AtXRN2/3. Additionally, two monocot XRNs were identified that are only distantly related to the Arabidopsis XRNs.

**Figure 1.**
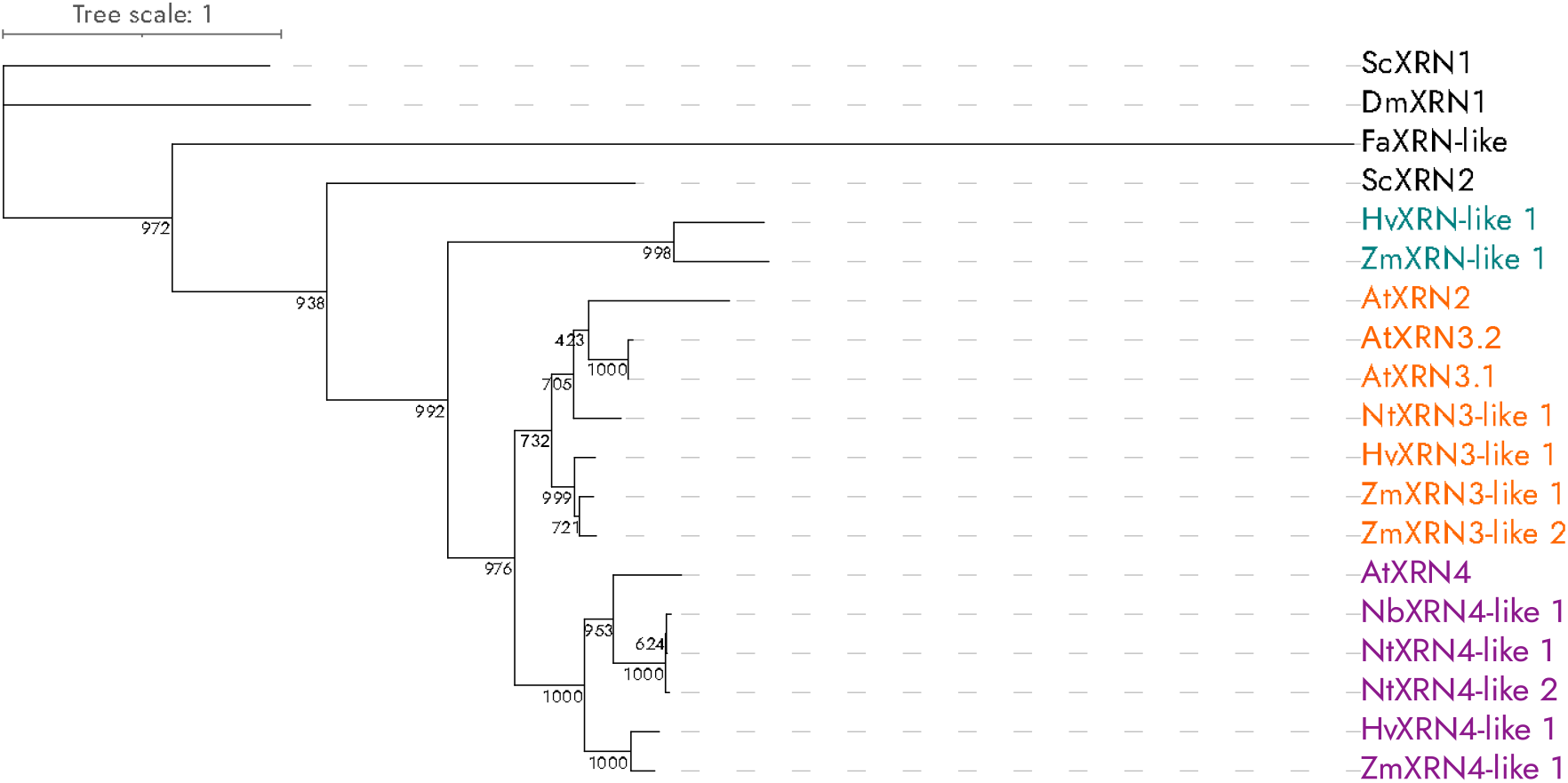
Phylogenetic analysis reveals three distinct plant XRN clades. The maximum-likelihood-based tree was generated from a multiple sequence alignment of XRN protein sequences identified by BLAST. The plant XRNs cluster into a XRN4-like clade (purple), a XRN2/3-like clade (orange), and an outgroup comprising two monocot enzyme sequences (teal). Numbers on nodes indicate the respective bootstrap values from 1,000 replicates. Branch lengths represent phylogenetic distances. Dm: *D. melanogaster*, Sc: *S. cerevisiae*, Fa: *F. algeromassiliense*, At: *A. thaliana*, Nt: *N. tabacum*, Nb: *N. benthamiana*, Hv: *H. vulgare*, Zm: *Z. mays*.

### For twelve of fifteen plant XRNs, the conserved active center was predicted with ssRNA in 5′ direction *in silico*

The crucial and diverse purposes of the XRN family are likely driven by the specific structure of these enzymes. Currently, no crystal structures of plant XRNs, either alone or in complex with a nucleotide substrate, are available. To gain a better understanding about their mechanistic function, computational structure prediction of the protein sequences with 5 nt long ssRNA and two magnesium ions was performed via Chai-1. Plausible models were selected by criteria of the active center known in the literature: prominent tower domain helix, magnesium ions andssRNA in 5′ direction in the active center, and possibility of π-π stacking of 3 nts between the stacking His and stacking Trp [35,41,42]. Out of all fifteen plant XRN models, twelve were predicted to reproduce the conserved active center with ssRNA in 5′ direction (Figure 2, Figure 3 and Figure S 2). For NtXRN4-like 2 and FaXRN-like, plausible models were obtained by structure prediction without contact restraints or template. With the crystal structure of ScXRN2 defined as template, plausible models were obtained for ten enzymes, namely AtXRN4, NbXRN4-like 1, HvXRN4-like 1, ZmXRN4-like 1, AtXRN2, AtXRN3.1, AtXRN3.2, ZmXRN3-like 1, HvXRN-like 1, and ZmXRN-like 1. The model of NtXRN4-like 1 was obtained with contact restraints between the nt 1 and the stacking His, and the nt 3 and the stacking Trp. For NtXRN3-like 1, HvXRN3-like 1, and ZmXRN3-like 2, no plausible structures were obtained. Even with the mentioned modifications applied, the ssRNA was still predicted incorrectly in 3′ direction, although the ssRNA and the magnesium ions were localized in the active center and the tower domain helix was modeled correctly. Because no plausible models were generated, these three candidate enzymes were excluded from further analysis. The structure predictions show that conserved structure characteristics are recognized well in computational structure modeling.

**Figure 2.**
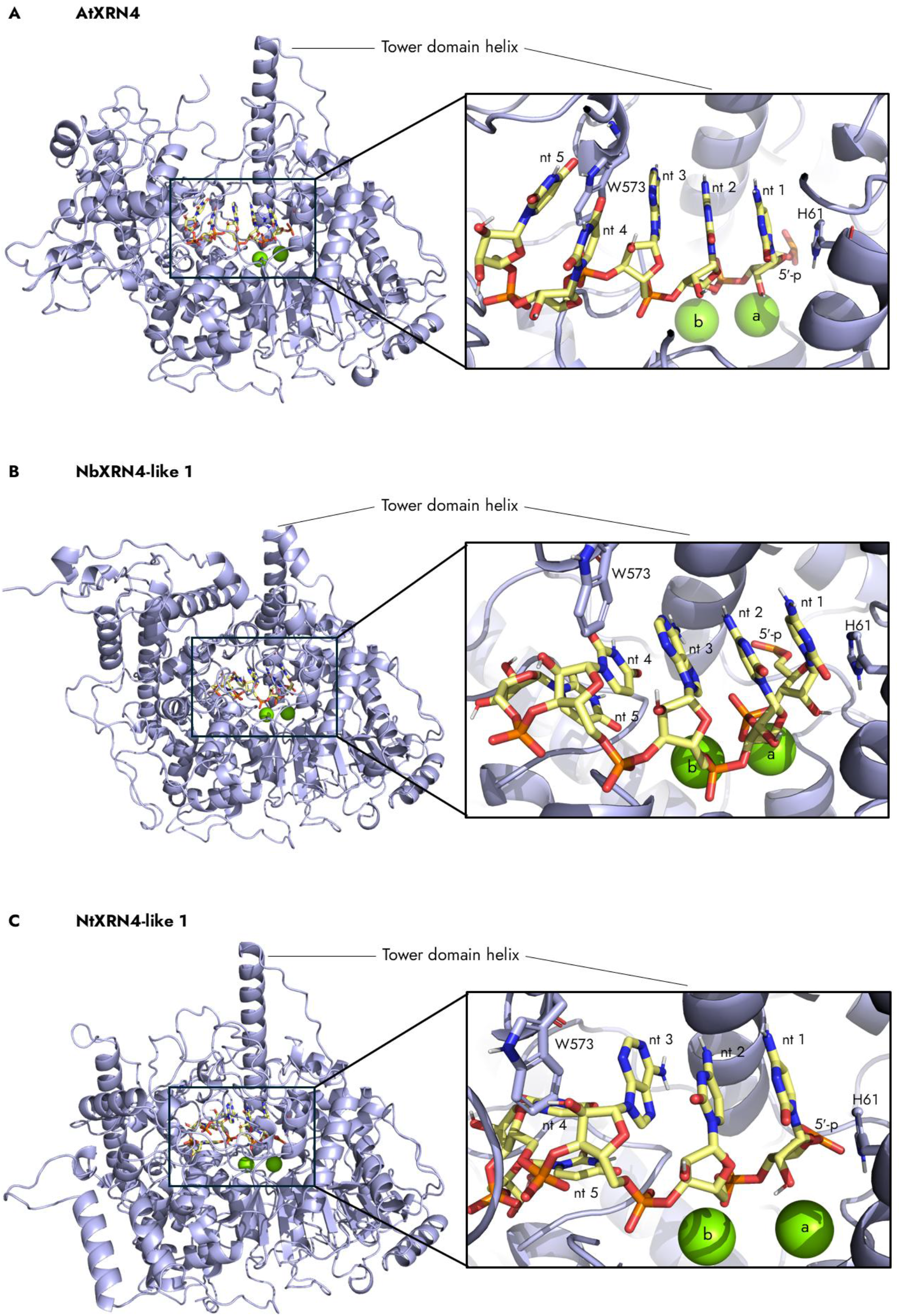
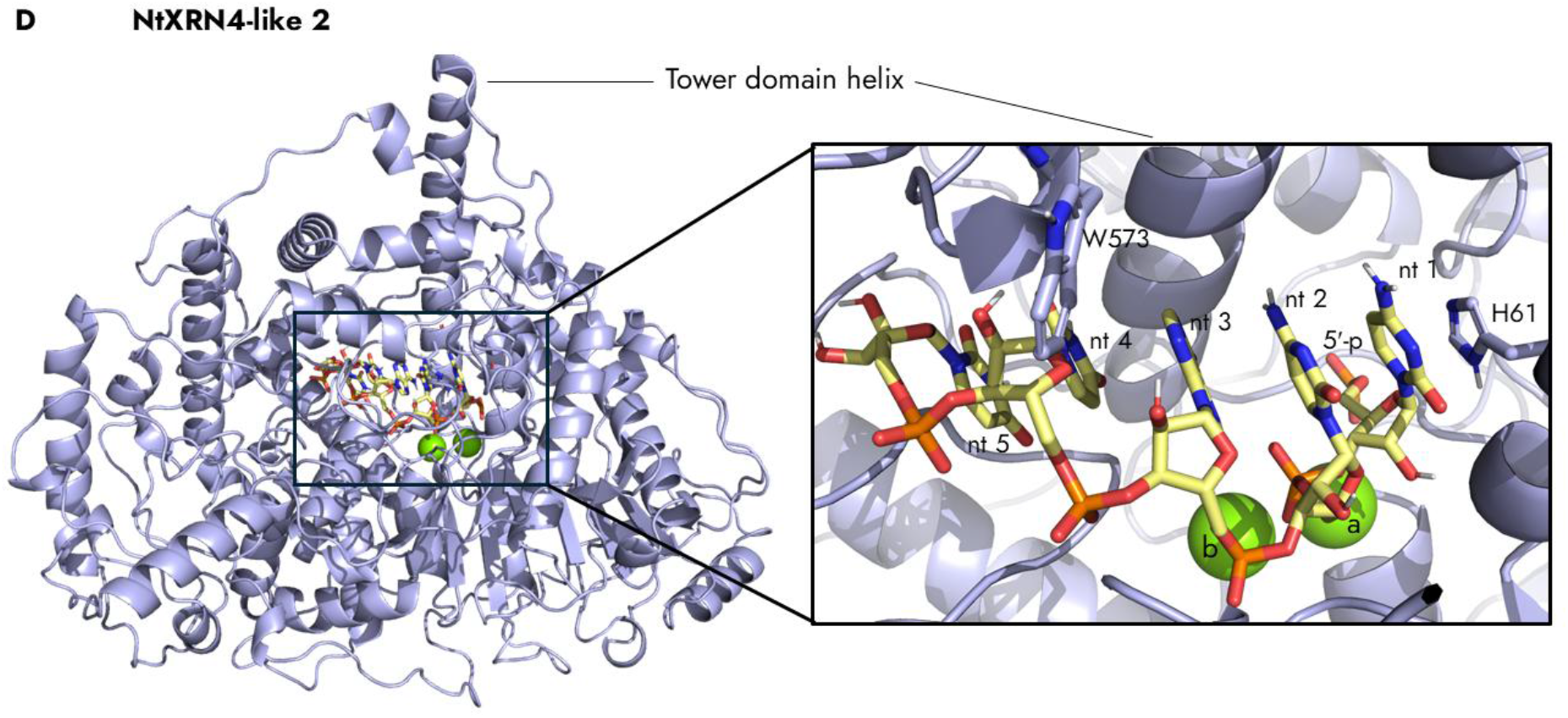
Structure predictions of the dicot members of the XRN4-like clade. The protein structures were predicted with a 5 nt long ssRNA and two magnesium ions (a, b) by Chai-1 and processed via MAXIMOBY. An overview of the enzyme structure (left) and a close-up of the active center (right) is shown for A. AtXRN4, B. NbXRN4-like 1, C. NtXRN4-like 1, and D. NtXRN4-like 2. Enzyme in light blue, ssRNA in yellow, atoms colored by element: hydrogen in white, nitrogen in blue, oxygen in red, sulfur in orange, and magnesium in green. W: tryptophan, H: histidine, 5′-p: 5′-monophosphate, nt: nucleotide.

**Figure 3.**
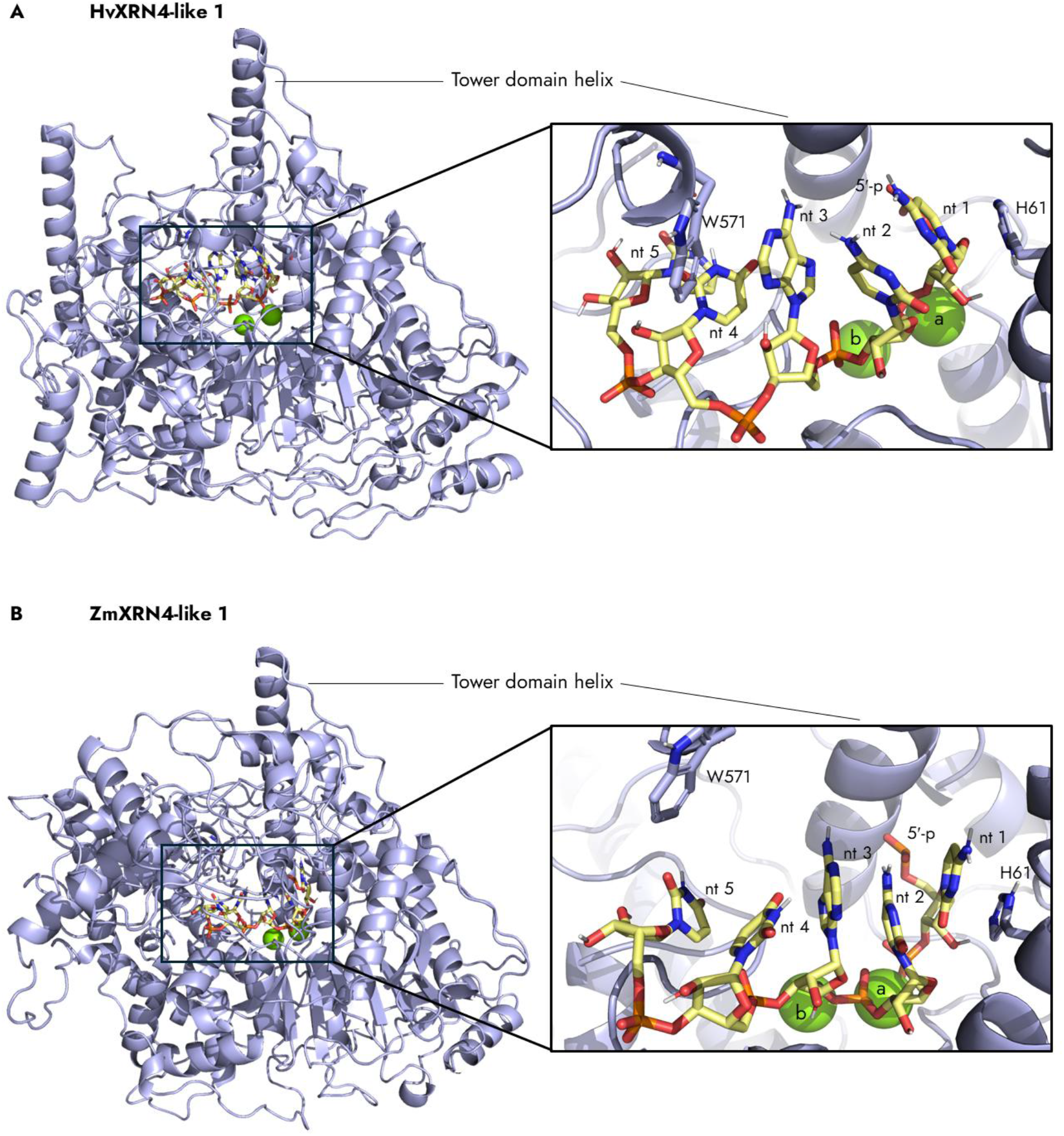
Structure predictions of the monocot members of the XRN4-like clade. The protein structures were predicted with a 5 nt long ssRNA and two magnesium ions (a, b) by Chai-1 and processed via MAXIMOBY. An overview of the enzyme structure (left) and a close-up of the active center (right) is shown for A. HvXRN4-like 1, and B. ZmXRN4-like 1. Enzyme in light blue, ssRNA in yellow, atoms colored by element: hydrogen in white, nitrogen in blue, oxygen in red, sulfur in orange, and magnesium in green. W: tryptophan, H: histidine, 5′-p: 5′-monophosphate, nt: nucleotide.

## In eleven of twelve processed plant XRN models, four distinct amino acids within the active center interact with nt 1 and the magnesium ions

In the active center of the homologous XRNs, the RNA substrate is stabilized by non-covalent interactions with amino acids of the active center, such as the π-π stacking interactions involving His and Trp, as well as by direct and indirect coordination of the magnesium ions [35,41,42]. To identify amino acids potentially involved in ssRNA stabilization and magnesium coordination, contacts between amino acids, nts, and magnesium ions were analyzed in the energy-optimized structure prediction models of the plant XRNs. The magnesium ions were designated as magnesium ion a and b according to their order in the 5′ direction of the RNA substrate. Here, we focused on direct contacts of the magnesium ions with amino acids and nts, as well as aromatic interactions between amino acids and nts. Attention was paid to centered and edge ring-ring contacts, with centered contacts being especially relevant for π-π stacking, while edge contacts could also contribute to π-π stacking in a dynamic enzyme. In all twelve plant XRN models, the stacking His interacted with nt 1, while Glu and two structurally neighboring Asp residues interacted with the magnesium ions in eleven of twelve plant XRN models (Figure 4). The magnesium ions of the plant XRN models were exclusively coordinated by negatively charged Asp and Glu residues (Figure 4 A, B). Three amino acids interacted with the magnesium ions in eleven of twelve plant XRN models: Glu (in DmXRN1: Glu177 [35]) and the two structurally neighboring Asp residues (in DmXRN1: Asp205, Asp207 [35]) (Table 1). All interactions of the second of the two structurally neighboring Asp residues involved magnesium ion b, with two of the eleven models predicting interactions with both magnesium ions. The magnesium ions also interacted with another Asp (in DmXRN1: Asp288 [35]) in eight models and with the Asp of the tower domain helix (in DmXRN1: Asp86 [35]) in seven models. Additionally, the magnesium ions interacted with nt 2 six times, with nt 1 thrice, and with nt 3 twice. In all twelve plant XRN models, the stacking His (in DmXRN1: His41 [35]) interacted with nt 1 via centered and edge ring-ring interactions (Figure 4 C, D). Out of these, centered ring-ring interactions were found for the models NtXRN4-like 2, AtXRN3.1, ZmXRN3-like 1, and ZmXRN-like 1. In six of the plant XRN models, the stacking Trp (in DmXRN1: Trp540 [35]) interacted with nt 3 via edge ring-ring interactions, with an additional centered ring-ring interaction in the AtXRN4 model (Figure 4 E, F). In addition, the stacking Trp of the AtXRN3.1 model interacted with nt 4 via an edge ring-ring interaction and both backbones of nt 4 and the stacking Trp of the NtXRN4-like 2 model formed a hydrogen bond (Figure S 3 C, D). A total of eight models showed interactions involving the stacking Trp. Arginine (Arg805, not conserved) showed centered ring-ring interactions with nt 2 and nt 3 in model HvXRN-like 1 (Figure 4 F and Figure S 3 B). Several additional amino acids interacting with the ssRNA in the plant XRN models were identified and further analyzed in the sequence motif analysis. In summary, in all twelve analyzed models, contacts were identified for magnesium ion b, nt 1 and nt 5 (Figure 4 A-D and Figure S 3 E, F). Contacts involving magnesium ion a and nt 3 were found in eleven and in ten models, respectively (Figure 4 A, B, E, F). Contacts involving nt 2 and nt 4 were found in six and in seven models, respectively (Figure S 3 A-D). These findings confirm the previously known amino acids involved in magnesium coordination and π-π stacking and indicate that these interactions are also relevant in plant XRNs.

**Figure 4.**
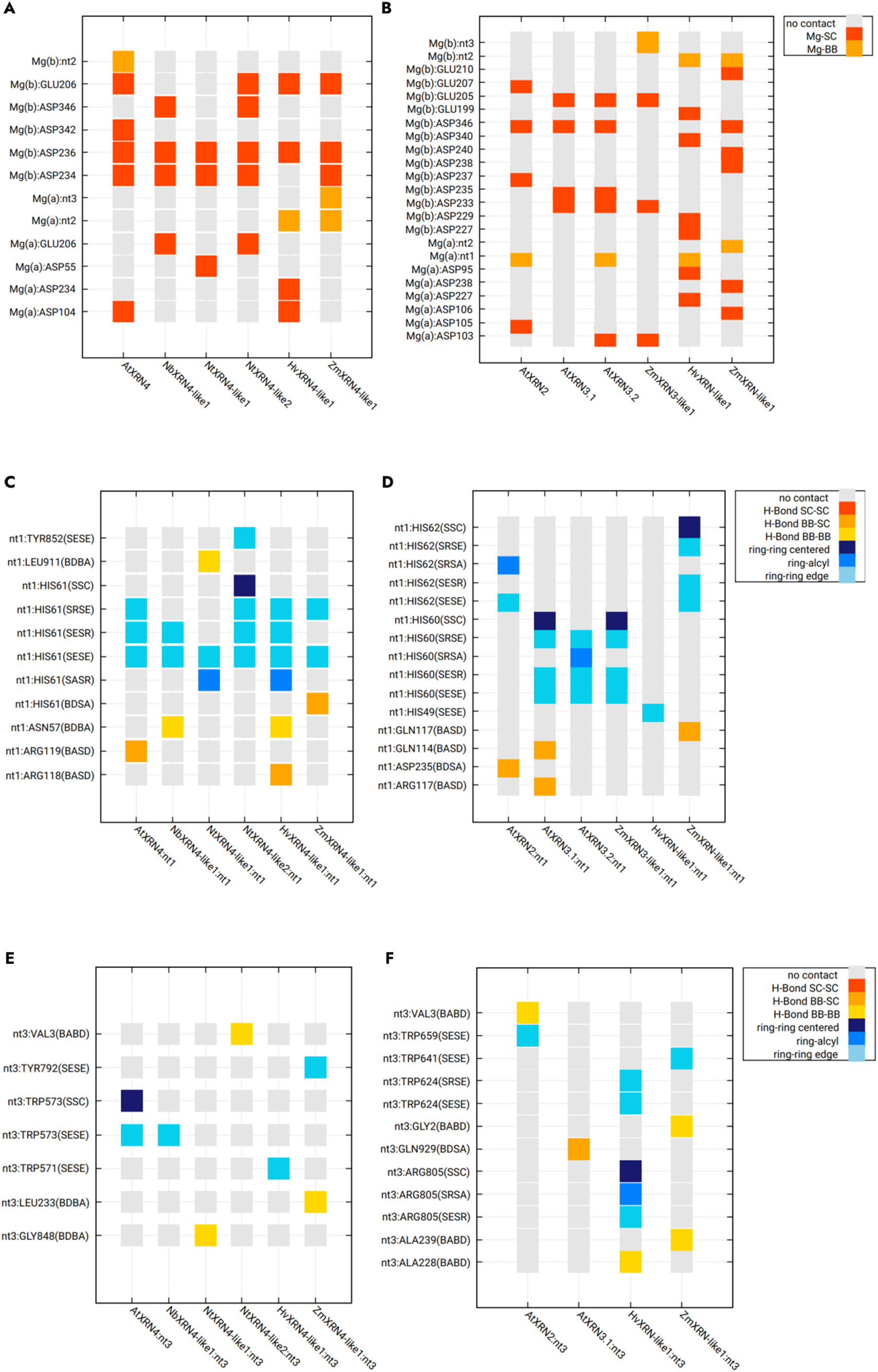
Contact analysis via kontakteUR revealed amino acids that participate in the mediation of magnesium ions and ssRNA. A, B. Contacts of the magnesium (Mg) ions with A. the XRN4-like clade models and B. the models of the XRN2/3-like clade and the monocot outgroup. C, D. Contacts of nt 1 with C. the XRN4-like clade models and D. the models of the XRN2/3-like clade and the monocot outgroup. E, F. Contacts of nt 3 with E. the XRN4-like clade models and F. the models of the XRN2/3-like clade and the monocot outgroup. SC in legend: side chain, BB in legend: backbone, Abbreviations of the y-axis: S: side chain, B: backbone. In hydrogen bonds: H: hydrogen, D: H-donor, A: H-acceptor. In ring interactions: R: ring (face), E: ring (edge), A: alkyl group, C: ring (center). nt: nucleotide.

**Table 1.** The seven identified conserved sequence motifs and the stacking Trp position within the XRN catalytic domain were compared with reported important amino acids and their function.

| Designated motif name | Known structure | DmXRN1 | ScXRN1 | ScXRN2 | AtXRN4 | Reported function | Contacts in this study |
|---|---|---|---|---|---|---|---|
| N-terminal motif: MGXPXF | Comprises the N-terminal α1 [35] | 1-6 aa | 1-6 aa | 1-6 aa | 1-6 aa | N-terminal α1 forms a narrow channel for substrate entry site [35] | Interaction of Met, Gly, Pro, and Phe with nt 2, nt 3 and nt 5 |
| Stacking His motif: YLDXNX(3)H | Comprises both β2 and α2 partly [35] | 33-41 aa | 33-41 aa | 51-59 aa | 53-61 aa | His for substrate hydrolysis and nucleotide stacking, Asp and Asn for Mg<sup>2+</sup> coordination [35,41,42] | Interaction of the stacking His and Asn with nt 1, as well as one interaction of Asp with magnesium ion a |
|  |  | Asp35 [35] | Asp35 [41] | Asp54 | Asp55 |  |  |
|  |  | Asn37 | Asn37 [41] | Asn55 | Asn57 |  |  |
|  |  | His41 [35] | His41 [41] | His59 [42] | His61 |  |  |
| Tower domain helix: DGVAPXAKMXQQRXRR | Tower domain helix, α4 [35] | 86-101 aa | 86-101 aa | 102-117 aa | 104-119 aa | Asp for Mg<sup>2+</sup> coordination and Lys, Gln and both Arg residues form the 5′-p basic pocket [35,41,42] | Asp coordinated exclusively magnesium ion a, Lys interacted with nt 2, and the second Gln and the last both Arg residues interacted with nt 1. |
|  |  | Asp86 [35] | Asp86 [41] | Asp102 | Asp104 |  |  |
|  |  | Lys93 [35] | Lys93 | Lys109 [42] | Lys111 |  |  |
|  |  | Gln97 [35] | Gln97 | Gln113 [42] | Gln115 |  |  |
|  |  | Arg100-101 [35] | Arg100-101 | Arg116-117 [42] | Arg118-119 |  |  |
| GEGE motif: GEGEHKX(4)I | α6 [35] | 174-184 aa | 175-185 aa | 202-212 aa | 203-213 aa | Both Glu residues for Mg<sup>2+</sup> coordination [35,41,42] | The second Glu coordinated magnesium ion b, magnesium ion a or both at once. |
|  |  | Glu175 | Glu176 [41] | Glu203 | Glu204 |  |  |
|  |  | Glu177 [35] | Glu178 [41] | Glu205 [42] | Glu206 |  |  |
| DAD motif: YGLDADLIXLXLX(9)LRE | α7 and β6 [35] | 202-225 aa | 203-226 aa | 230-253 aa | 231-254 aa | Both Asp residues for Mg<sup>2+</sup> coordination [35,41]. | Both Asp residues for Mg<sup>2+</sup> coordination, predominantly of magnesium ion b. Additionally, interaction of the first Leu, the first Asp, and Ala with nt 1 to 4. |
|  |  | Asp205 [35] | Asp206 [41] | Asp233 [42] | Asp234 |  |  |
|  |  | Asp207 [35] | Asp208 [41] | Asp235 [42] | Asp236 |  |  |
| GNDF motif: GNDFXPXXP | / | 286-294 aa | 289-297 aa | 331-339 aa | 340-348 aa | In DmXRN1, Asp288 is predicted to coordinate the non-resolved Mg<sup>2+</sup> [35] | Interactions exclusively with magnesium ion b and nt 5 |
|  |  | Asp288 [35] | Asp291 | Asp333 | Asp342 |  |  |
| YY motif: YYX(5)SWXWXY | Partially α13 [35] | 449-461 aa | 547-559 aa | 641-653 aa | 482-494 aa | / | Highest motif score of all but no interactions with ssRNA nor Mg<sup>2+</sup> |
| Stacking Trp position: DX(3)KX(3)[WH] | Trp540-containing loop [35] | 532-540 aa | 630-638 aa | 724-732 aa | 565-573 aa | Loop forms a narrow channel for substrate entry site and stacking Trp for nucleotide stacking [35,41,42] | Interaction of the stacking Trp with nt 3 and nt 4 as well as interaction of the conserved Lys with nt 5 |
|  |  | Trp540 [35] | Trp638 [41] | Trp732 [42] | Trp573 |  |  |

### Seven potentially functional motifs were identified within the protein sequences

Homologous proteins usually contain conserved amino acids within functionally distinct domains, which can serve as targets for mutagenesis studies. These conserved amino acids can be detected by aligning the protein sequences of these homologous proteins. Further information on enzyme function and subcellular localization can be derived from sequence analysis. To identify conserved amino acid motifs within the active center of the plant XRNs that are involved in the mediation of ssRNA and magnesium ions, a combination of the previous structure-based contact analysis and the sequence motif analysis was utilized. All XRN protein sequences were subjected to GLAM2 to predict conserved sequence motifs and an MSA was generated using Clustal Omega. Seven conserved motifs were identified: N-terminal motif, stacking His motif, Tower domain helix, GEGE motif, DAD motif, GNDF motif, and YY motif (Figure 5 A-C, Figure S 4 and Figure S 6). Five of these seven motifs were identified by GLAM2 and numbered from the N-terminus, although three of them were only partially identified compared to the MSA and the structure-based contact analysis (Figure 5 B, C and Figure S 4). The N-terminal motif started at the first amino acids of all sequences. In the analyzed plant XRN models, this motif had a total of eleven contacts with nt 2 to nt 5, eight of which were with nt 5 (Figure S 6). Two additional contacts with nt 5 were found for two non-conserved amino acids directly following this motif. The predominance of contacts between the N-terminal motif and nt 5 suggests a potential function involving nt 5. The stacking His motif was named after the stacking His and corresponded to GLAM2 motif 1. For this motif, GLAM2 did not include the FaXRN-like sequence in the alignment (Figure S 4 A). To highlight the most important amino acids of all protein sequences, an adjusted version of this motif containing only universally conserved amino acids is shown (Figure 5 A). In addition to the previously mentioned contacts between the nt 1 and the stacking His in all plant XRN models, the stacking His motif formed two contacts with nt 1 via Asn and one contact with magnesium ion a via Asp (Figure S 6). The contacts of the stacking His, Asn, and Asp with nt 1 or with magnesium ion a next to nt 1 demonstrate the function of the stacking His motif at the 5′ end of the ssRNA. GLAM2 motif 2 identified the tower domain helix, which overall showed the second highest score among all motifs predicted by GLAM2 (Figure 5 B). In addition to the seven interactions of Asp with magnesium ion a, the Tower helix domain interacted with nt 1 twice via the second glutamine, thrice via the neighboring arginines and four times with nt 2 via lysine (Figure S 6). These findings suggest that the tower domain helix interacts mainly with magnesium ion a and less-frequently with nt 1 and nt 2. The GEGE motif was derived from the MSA and contact matrix analysis. The second Glu interacted with the magnesium ions in eleven plant XRN models, with ten contacts involving magnesium ion b and two contacts involving magnesium ion a, as NtXRN4-like 2 contacted with both magnesium via Glu (Figure S 6). Notably, however, this motif was not identified by GLAM2. These findings suggest a predominant function of the GEGE motif in the coordination of magnesium ion b. The DAD motif was identified partially as GLAM2 motif 3 and had the lowest overall score of all GLAM2 motifs (Figure S 4 B). As described earlier, the two structurally neighboring Asp residues interacted with the magnesium ions in eleven plant XRN models. In nine models, both Asp residues interacted with magnesium ion b simultaneously. The first Asp also interacted once with nt 1.

**Figure 5.**
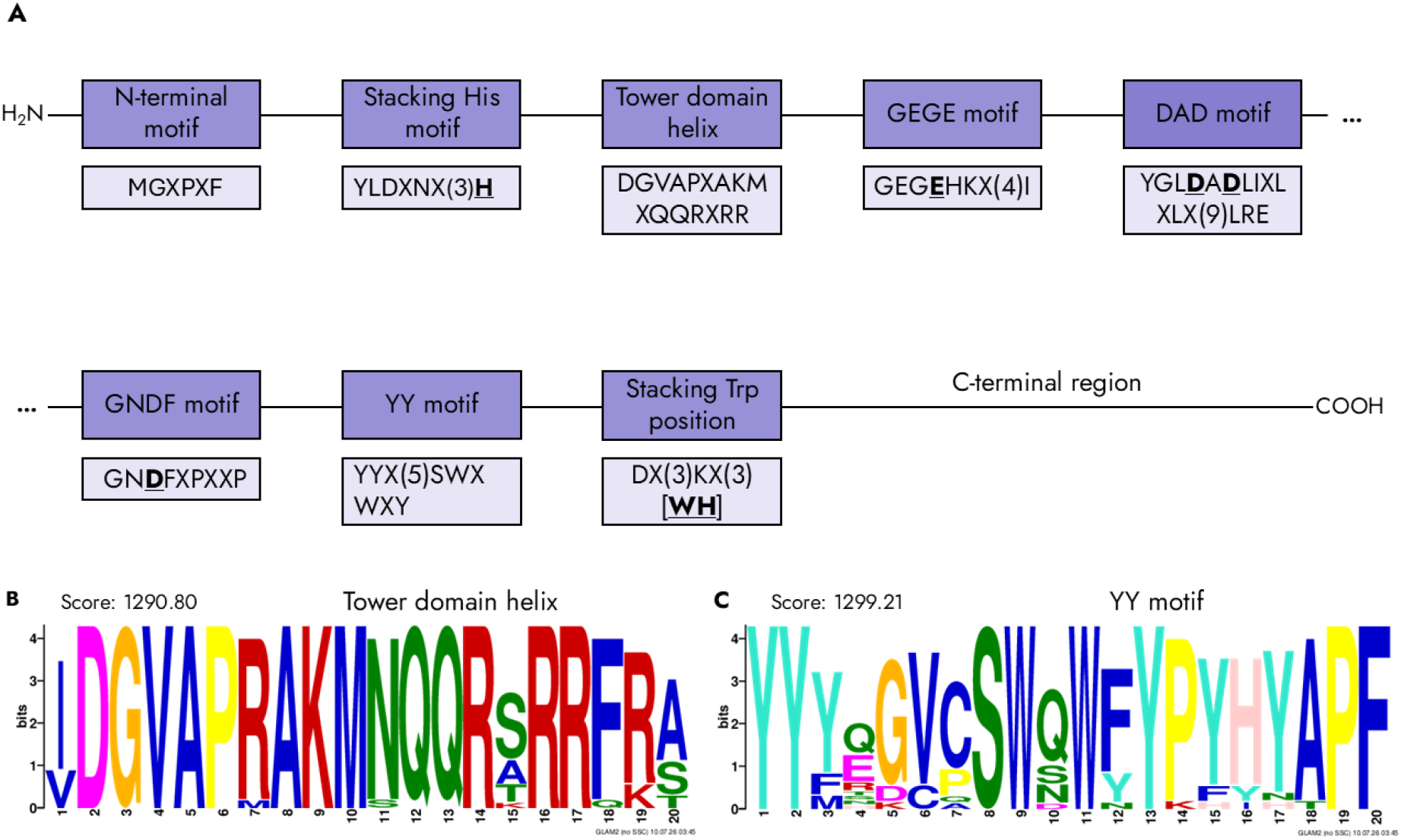
Sequence motif analysis shows seven conserved motifs. A. Schematic overview of the localization and distribution of conserved motifs (violet) within the investigated protein sequence of XRN family members. Conserved amino acids within each motif are indicated below (light violet). The amino acid motifs show conserved amino acids throughout all sequences in 1-letter code. X indicates variable amino acid residues (when one amino acid was different). Exception was for [WH] in the stacking Trp position. B, C. Motifs completely identified by GLAM2 visualized as sequence logo plot with B. showing the tower domain helix logo plot and with C. showing the YY motif logo plot.

Additionally, the first leucine interacted once each with nt 1 and nt 2, while the alanine between the two Asp residues interacted thrice with nt 3 and nt 4 (Figure S 6). This indicates that the DAD motif predominantly interacts with magnesium ion b. The GNDF motif contained the Asp, which was shown to interact with magnesium ion b in eight plant XRN models in the contact analysis. This motif was identified as GLAM2 motif 4 with the second-lowest score of all motifs, although the initial glycine was missing (Figure S 4 C). In the region of the GNDF motif, eight additional interactions with nt 5 were found (Figure S 6). This suggests that the GNDF motif interacts with magnesium ion b and nt 5. The YY motif was identified as GLAM2 motif 5 and had the highest score of all motifs (Figure 5 C). However, no interactions of the plant XRN models with neither ssRNA nor magnesium ions were found in the structure-based contact analysis (Figure S 6). This suggests that the YY motif is conserved, but the functional relevance remains elusive. The stacking Trp was located downstream of all identified motifs and was not considered conserved in the MSA, as the FaXRN-like sequence contained a His at this position (Figure S 6). In the predicted FaXRN-like structure model, this His was positioned in a manner consistent with a potential π-π stacking interaction. Therefore, this position might still be of functional relevance, as His can also mediate π-π stacking interactions. Thus, the aromatic character of this position could be considered conserved. To evaluate the potential secretion of the plant XRNs into the apoplast, the protein sequences of the plant XRN candidates were screened for the presence of a signal peptide motif via SignalP. For the plant XRN, ScXRN1, ScXRN2 and DmXRN1 sequences submitted to SignalP, no secretory signal peptide was identified that would mediate the transport to the apoplast (Figure S 5). This indicates the absence of a classical secretory signal peptide in the plant XRNs. Overall, this integrated analysis identified seven conserved motifs and several conserved amino acids in the plant XRN candidates, which could serve as important targets for future mutational studies.

### xrRNA-XRN complex modeling - structural volume of the xrRNA model compared to NbXRN4-like 1 model

Research on different viral xrRNA motifs has shown that they can protect downstream areas of viral RNAs from degradation by host XRNs [44]. Therefore, we hypothesized that the incorporation of the xrRNA motif into the sequences of sprayable RNAs could potentially also protect ssRNA from degradation during spray applications. To compare the structural volume of the ST9a xrRNA crystal structure [44] with that of the plant XRNs, a ST9a xrRNA-tagged ssRNA sequence was used for an AlphaFold3 structure prediction, the predicted xrRNA sequence was then replaced by the ST9a xrRNA crystal structure, and the resulting 3D xrRNA model (Figure 6 A) was then modeled in complex with the NbXRN4-like 1 structure prediction model (Figure 6 B). As a first xrRNA-XRN complex model suggestion, the NbXRN4-like 1 model was chosen because *N. benthamiana* can serve as a host for ST9a infection in experimental setups [44]. The second nucleotides of both the tagged-RNA model and the ssRNA of the NbXRN4-like 1 model were aligned to model the potential degradation complex of the 3D xrRNA model by the enzyme. This revealed that the 3D xrRNA model occupies a structural volume much larger than the substrate entry site of the NbXRN4-like 1 model (Figure 6 C). This finding suggests that the xrRNA fold could also sterically hinder the entry of the tagged RNA into the plant XRNs. This static model of a dynamic complex builds a basis for future molecular dynamics simulation to test the xrRNA fold stability in a physiologic solution *in silico*. Furthermore, to test whether the xrRNA motif increases the resistance of sprayed mRNA in future analyses, the xrRNA of ST9a together with an upstream leader sequence was generated by IVT and modeled *in silico*. Two different versions of the xrRNA motif were generated: one was unmodified, while the second version included two point mutations (A12U/A13U) in the L2A region, which is expected to interfere with the correct formation of the protective ring structure of the xrRNA motif. The IVT products were analyzed on a 6% urea polyacrylamide gel, which showed the expected bands at a size of 114 nt (Figure 6 D and Figure S 7 A). This implies that the IVT of xrRNA-tagged RNA was successful and that these ssRNAs can be used for *in vitro* RNA degradation assays with plant XRNs in the future.

**Figure 6.**
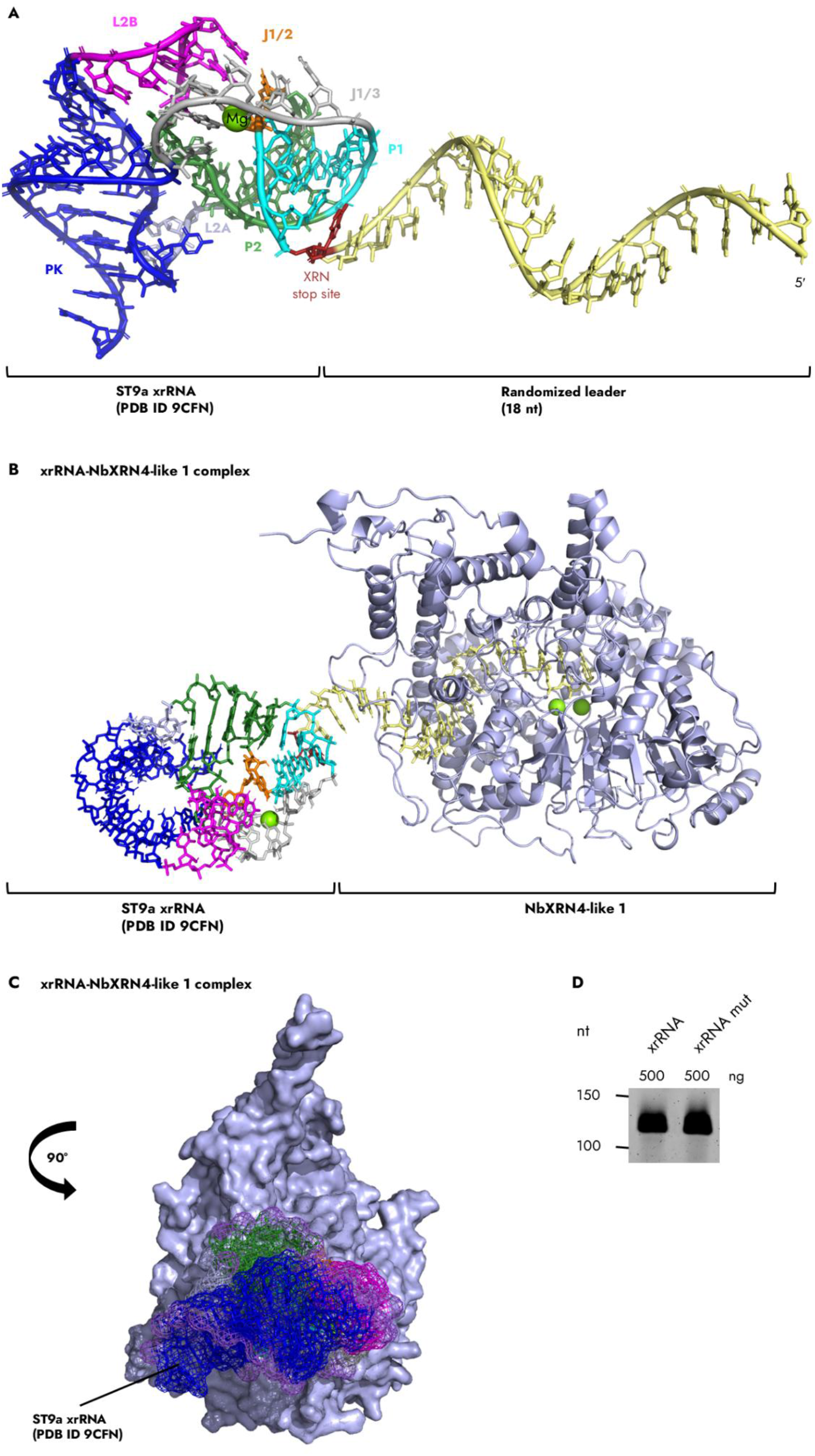
xrRNA-XRN complex modeling reveals structural volume of the xrRNA model compared to NbXRN4-like 1 model. A. xrRNA model with leader (yellow) and ST9a xrRNA crystal structure (PDB 9CFN). Mg: magnesium ion, 5′: 5′ end. B. xrRNA model in complex with NbXRN4-like 1 model (light blue). C. Predicted structural volume of xrRNA model and NbXRN4-like 1 model. D. 6% urea polyacrylamide gel showing the xrRNA design without and with two point mutations (A12U/A13U) in the L2A region at its expected size of 114 nt. mut: mutated, nt: nucleotide, ng: nanograms. Different regions of ST9a xrRNA colored in cyan (P1), orange (J1/2), green (P2), light blue (L2A), blue (PK), magenta (L2B), grey (J1/3).

## Discussion

The aim of this study was the computational sequence- and structure-based analysis of the XRN family in the plant species Arabidopsis, maize, barley, *N. benthamiana* and *N. tabacum*. The members of XRN family in plants were characterized in more detail using *in silico* structure modelling. Phylogenetic analysis of the plant XRN candidates identified by BLASTP along with DmXRN1, ScXRN1/2, and FaXRN-like revealed two distinct XRN clades in plants, the XRN4-like clade including AtXRN4, the XRN2/3-like clade including AtXRN2 as well as both isoforms of AtXRN3, and a separate outgroup comprising two monocot XRN-like sequences. The phylogenetic relationships were supported by high bootstrap values. This phylogenetic division aligns with previous phylogenetic analysis, where ScXRN1 was shown to be phylogenetically more distant to the Arabidopsis XRNs compared to ScXRN2. [21]. Here, also AtXRN2 and AtXRN3 where shown to cluster together [21]. This agreement supports the reliability of the phylogenetic tree established here. Notably, this tree included XRNs from plant species for which comparatively little phylogenetic information is available, including barley, maize, and tobacco. Furthermore, the outgroup comprising two monocot XRNs raises the question of whether this outgroup is restricted to monocots and whether its members fulfil a distinct function. Future studies should therefore focus on identifying further members of this outgroup and investigating their cellular localization and potential functions. The high similarity of ScXRN2 to the plant XRNs analyzed here also makes it a valuable model for future structural and functional studies when experiments with plant XRNs are challenging.

The protein sequences of the plant XRN candidates and the xrRNA design were submitted to AI-based structure prediction tools, such as Chai-1 and AlphaFold3, that use trained neural networking. Twelve of fifteen predicted plant XRN models showed the conserved active center with ssRNA in 5′ direction according to the criteria found in the literature [35,41,42]. Template or contact restraints were used for eleven of the twelve plant XRN models. Furthermore, all fifteen plant XRN models predicted the specific active center with the two magnesium ions and the ssRNA; although, for three of the fifteen models, the ssRNA was incorrectly placed in 3′ direction. Additionally, for computational RNA structure prediction, the 3D structure of the xrRNA design was predicted seemingly close to the crystal structure of ST9a xrRNA. These findings support the current knowledge on AI-based structure prediction that single protein structure predictions resulting from deep learning are more accurate, as they outperform *de novo* protein structure predictions without deep learning techniques [78–82]. However, the accuracy is slightly decreased for protein complex models in the interface area and is also limited for predicted RNA structure models [78,79,83]. Thus, combining AI-based structure prediction tools and homologous templates of experimentally determined crystal structures appears to improve the accuracy of structure prediction for proteins in complex with RNA. Insights generated from this combined modeling approach are applicable to generate more accurate structure predictions for other homologous proteins in complex with RNA where no crystal structures are available

In eleven of twelve plant XRN models, four distinct amino acids within the active center were identified that interact with nt 1 and two magnesium ions. They and several other interacting amino acids were found to be conserved within six potentially functional motifs in the N-terminal half of all aligned protein sequences. This region aligned with the XRN catalytic domain of DmXRN1 (1-674aa) and the catalytic domain of ScXRN1 (1-772aa) [35,41]. The amino acids of the motifs named N-terminal motif, stacking His motif, the tower domain helix, GEGE motif, DAD motif, GNDF motif participated in ssRNA and magnesium ion mediation in the analyzed XRN models. Other less conserved amino acids of the N-terminal motif are suggested to be part of a positively charged rim of the active center located in proximity to 5′-p of nt 4 [35]. most contacts of the N-terminal motif involved nt 5. In previous mutation assays of ScXRN1 and ScXRN2, mutations of several amino acids affect exonuclease activity: the conserved Asn and the stacking His of the respective motif, the conserved Asp, glycine, lysine, the second glutamine and the second arginine of the tower helix domain motif, both Glu residues of the GEGE motif, and the two structurally neighboring Asp residues of the DAD motif [38,40,84]. All the mentioned conserved amino acids except the glycine of the tower domain helix motif and the first Glu of the GEGE motif were also found to interact in all plant XRN models. In ScXRN1 and partially in DmXRN1, the magnesium ions are mediated by the conserved amino acids Asp and Asn of the stacking His motif, Asp of the tower domain helix motif, the two conserved Glu residues of the GEGE motif, and the two Asps of the DAD motif [35,41]. Additionally, the tower domain helix motif consists of the conserved amino acids of the positively charged 5′-p binding pocket in DmXRN1 [35]. In the models generated in this work, the GNDF motif interacted with nt 5 and magnesium ion b. The YY motif found by sequence motif analysis with the highest score was the only motif, where no contacts between its amino acids and the ssRNA or the magnesium ions were identified. This contradiction suggests an allosteric function and requires further investigation. For both GNDF and YY motif, no suggestions about their function were found in the literature. In general, all plant XRN models are computational estimates and illustrate static structures. The variable contact points between plant XRN models might picture conformational changes such as shown for a member of the XRN family from *Chaetomium thermophilum in vitro* [85]. Ultimately, the findings of this work remain to be validated by experimental setups such as structure biology methods, mutagenesis studies or protein-RNA interaction assays. Further, molecular dynamics simulations would be a valuable tool to capture dynamic contacts as well as allosteric and conformational changes, and to assess the stability of the predicted models.

Overall, this computational analysis has revealed six potentially functional motifs with conserved amino acids that seem to be important for ssRNA and magnesium ion mediation, as well as one additional conserved motif with a potential allosteric function In summary, this study combined sequence and structural analyses to confirm previously reported conserved amino acids and structural domains (Mg^2+^-coordinating amino acids, stacking His and Trp, N-terminal α1, Tower domain helix, Trp540-containing loop), extend them to plant XRNs and identify conserved sequence motifs (N-terminal motif, Stacking His motif, Tower domain helix motif, GEGE motif, DAD motif, GNDF motif, YY motif). The integration of structural information and contact analysis enabled a detailed characterization of six of the conserved motifs, including the potential functional role of its conserved residues. These conserved residues and motifs are present in all XRNs analyzed here, making them promising targets for future mutagenesis studies. Ultimately, these insights may serve as a basis for identifying functionally relevant motifs in other RNA-cleaving enzymes.

Since sprayed ssRNAs need to pass the apoplast before entrance into the cell, these ssRNAs would be susceptible to degradation by apoplastic RNases. Previously described examples of such apoplastic RNases include 3′–5′ exoribonucleases of the RNase T2 family [86]. These RNases rely on the conventional secretory pathway, which is triggered by the presence of a signal peptide [86]. To test whether one of the identified XRN candidates might also be trafficked into the apoplast, the protein sequences of all plant XRN candidates were screened for the presence of a signal peptide. However, no secretory signal peptide motif was found in none of the candidates, which suggests that these plant XRN candidates probably are not secreted to the apoplast. On the other hand, unconventional protein secretion is still possible for the plant XRNs [87,88]. To investigate this possibility, a microscopic analysis of GFP-tagged XRNs should be performed for visualization of their localization *in planta*. However, XRNs especially the cytoplasmic XRN4-like clade form a cytoplasmic barrier after cellular uptake.

One possibility to protect ssRNAs from degradation by XRNs is the inclusion of the xrRNA motif into the sequence of sprayed ssRNAs. Previous research suggests that the xrRNA motif forms a ring-like tertiary structure that sterically hinders entrance into the XRN and by that inhibits degradation by *Kleuveromyces lactis* Xrn1 [44]. To evaluate transferability to plant XRNs and investigate the effect of the xrRNA motif in more detail, *in silico* modelling of an xrRNA-tagged RNA in complex with NbXRN4-like 1 has been performed. The resulting model indicated that the xrRNA possesses a structural volume that is too large to enter the modelled NbXRN4-like 1, indicating that the structure of the enzyme or the xrRNA would first have to unfold. Such process yields an energy barrier, hindering the degradation of RNA downstream of the xrRNA motif.

Classical MD simulations are well suited to assess the structural integrity and key tertiary contacts of the inhibitory xrRNA fold. Our xrRNA-XRN model serves as a base for such investigations. Moreover, to experimentally determine the function of the xrRNA motif, *in vitro* degradation assays on RNAs that have either been tagged with an unmodified xrRNA motif or mutated version of the xrRNA that is impaired in tertiary structure formation should be performed. To assess the function of xrRNA *in vivo*, a GFP-encoding mRNA should be tagged with the xrRNA motif and spray applied to plants, as the uptake of GFP-encoding mRNA was previously shown [11]. However, 5′-capped mRNA requires decapping or endonucleolytic cleavage before it can serve as a 5′-p substrate for XRNs. Furthermore, how, XRN stalling affects the cells and the whole plant needs further investigation, since plant XRNs have essential roles and malfunction as for AtXRN3 leads to embryo lethality [18,19].

Our approach shows that potentially functional amino acid motifs can be identified exclusively using computational methods, despite the above-discussed limitations In future studies we can expand our scope aiming for generalizability to other RNases.

## Conclusion

Here, we identified the potentially functional motifs of plant XRNs and their potential structural architectures, paving the way for further development in the computational recognition of other non-homologous RNA cleaving enzymes. The computational models predicted in this work could further support the rational design of exogenous ssRNAs with increased stability, since they allow to simulate the interaction of RNAs with XRNs *in silico* before starting more expensive wet lab experiments.

## Acknowledgements

We thank Moritz Dobrowitsch for providing the plasmids and for sharing his valuable knowledge and relevant literature which contributed to this project. We thank Dr. Veli-Vural Uslu for his help with identifying conserved motifs. We thank Dr. Timo Schlemmer for assisting in the creation of the graphical abstract.

## Funding

This work was supported by the Deutsche Forschungsgemeinschaft (DFG, SPP 2416, KO 4874/5-1), the Bavarian State Ministry of Food, Agriculture, Forestry and Tourism (StMELF) within the project “sweetRNA” (grant number 7346-1/139), and the European Union’s Horizon Europe research and innovation programme under the Marie Skłodowska-Curie grant agreement No. 101169421 (ViroiDoc).

Views and opinions expressed are those of the author(s) only and do not necessarily reflect those of the European Union or the European Research Executive Agency (REA). Neither the European Union nor the granting authority can be held responsible for them.

## Data availability

All data and supplemental materials used for the findings of this study are openly available in Zenodo via: https://doi.org/10.5281/zenodo.24546329

## References

[1] FAO, Plant Production and Protection, Food and Agriculture Organization of the United Nations (2026). https://www.fao.org/plant-production-protection/about/ (accessed April 28, 2026).

[2] M. Tudi, H. Daniel Ruan, L. Wang, J. Lyu, R. Sadler, D. Connell, C. Chu, D.T. Phung, Agriculture Development, Pesticide Application and Its Impact on the Environment, International Journal of Environmental Research and Public Health 18 (2021) 1112. 10.3390/ijerph18031112.

[3] W. Zhou, M. Li, V. Achal, A comprehensive review on environmental and human health impacts of chemical pesticide usage, Emerging Contaminants 11 (2025) 100410. 10.1016/j.emcon.2024.100410.

[4] J. Wesseler, The EU’s farm-to-fork strategy: An assessment from the perspective of agricultural economics, Applied Economic Perspectives and Policy 44 (2022) 1826–1843. 10.1002/aepp.13239.

[5] V. Silva, X. Yang, L. Fleskens, C.J. Ritsema, V. Geissen, Environmental and human health at risk – Scenarios to achieve the Farm to Fork 50% pesticide reduction goals, Environment International 165 (2022) 107296. 10.1016/j.envint.2022.107296.

[6] S.M. Hammond, S. Boettcher, A.A. Caudy, R. Kobayashi, G.J. Hannon, Argonaute2, a Link Between Genetic and Biochemical Analyses of RNAi, Science 293 (2001) 1146–1150. 10.1126/science.1064023.

[7] C. Rosa, Y.-W. Kuo, H. Wuriyanghan, B.W. Falk, RNA Interference Mechanisms and Applications in Plant Pathology, Annual Review of Phytopathology 56 (2018) 581–610. 10.1146/annurev-phyto-080417-050044.

[8] P. Barth, J. Drumm, A. Schmidt, F. Hartig, A. Koch, Efficiency of RNAi based gene silencing in fungi - a review and meta-analysis, (2026) 2026.03.04.709060. 10.64898/2026.03.04.709060.

[9] A. Chen, L. Halilovic, J.-H. Shay, A. Koch, N. Mitter, H. Jin, Improving RNA-based crop protection through nanotechnology and insights from cross-kingdom RNA trafficking, Current Opinion in Plant Biology 76 (2023) 102441. 10.1016/j.pbi.2023.102441.

[10] V.V. Uslu, A. Dalakouras, L. Sueselbeck, A. Koch, RNA Meets Agriculture: From Molecular Mechanisms to Market Applications, Annual Review of Plant Biology 77 (2026) 289–317. 10.1146/annurev-arplant-070825-083558.

[11] V.V. Uslu, M. Dobrowitsch, K.P. Danger, A.U. Furch, J. Noetzold, A.M. Richter, T. Schlemmer, Y. Dong, G. Krczal, A. Koch, Foliar mRNA spray induces protein synthesis in monocot crop and dicot model plant species, (2025) 2025.08.07.668951. 10.1101/2025.08.07.668951.

[12] G.C. MacIntosh, B. Castandet, Organellar and Secretory Ribonucleases: Major Players in Plant RNA Homeostasis, Plant Physiol 183 (2020) 1438–1452. 10.1104/pp.20.00076.

[13] C. Charbonnel, A. de Bures, J. Sáez-Vásquez, An In Vitro Approach To Study RNase III Activities of Plant RTL Proteins, in: M. Boudvillain (Ed.), RNA Remodeling Proteins: Methods and Protocols, Springer US, New York, NY, 2021: pp. 363–385. 10.1007/978-1-0716-0935-4_23.

[14] V.K. Nagarajan, C.I. Jones, S.F. Newbury, P.J. Green, XRN 5′ → 3′ exoribonucleases: Structure, mechanisms and functions, Biochimica et Biophysica Acta (BBA) - Gene Regulatory Mechanisms 1829 (2013) 590–603. 10.1016/j.bbagrm.2013.03.005.

[15] J. Houseley, D. Tollervey, The Many Pathways of RNA Degradation, Cell 136 (2009) 763–776. 10.1016/j.cell.2009.01.019.

[16] N.L. Garneau, J. Wilusz, C.J. Wilusz, The highways and byways of mRNA decay, Nat Rev Mol Cell Biol 8 (2007) 113–126. 10.1038/nrm2104.

[17] J.P. Kastenmayer, P.J. Green, Novel features of the XRN-family in Arabidopsis: Evidence that AtXRN4, one of several orthologs of nuclear Xrn2p/Rat1p, functions in the cytoplasm, Proceedings of the National Academy of Sciences 97 (2000) 13985–13990. 10.1073/pnas.97.25.13985.

[18] I. Gy, V. Gasciolli, D. Lauressergues, J.-B. Morel, J. Gombert, F. Proux, C. Proux, H. Vaucheret, A.C. Mallory, Arabidopsis FIERY1, XRN2, and XRN3 Are Endogenous RNA Silencing Suppressors, Plant Cell 19 (2007) 3451–3461. 10.1105/tpc.107.055319.

[19] K. Yamada, J. Lim, J.M. Dale, H. Chen, P. Shinn, C.J. Palm, A.M. Southwick, H.C. Wu, C. Kim, M. Nguyen, P. Pham, R. Cheuk, G. Karlin-Newmann, S.X. Liu, B. Lam, H. Sakano, T. Wu, G. Yu, M. Miranda, H.L. Quach, M. Tripp, C.H. Chang, J.M. Lee, M. Toriumi, M.M.H. Chan, C.C. Tang, C.S. Onodera, J.M. Deng, K. Akiyama, Y. Ansari, T. Arakawa, J. Banh, F. Banno, L. Bowser, S. Brooks, P. Carninci, Q. Chao, N. Choy, A. Enju, A.D. Goldsmith, M. Gurjal, N.F. Hansen, Y. Hayashizaki, C. Johnson-Hopson, V.W. Hsuan, K. Iida, M. Karnes, S. Khan, E. Koesema, J. Ishida, P.X. Jiang, T. Jones, J. Kawai, A. Kamiya, C. Meyers, M. Nakajima, M. Narusaka, M. Seki, T. Sakurai, M. Satou, R. Tamse, M. Vaysberg, E.K. Wallender, C. Wong, Y. Yamamura, S. Yuan, K. Shinozaki, R.W. Davis, A. Theologis, J.R. Ecker, Empirical Analysis of Transcriptional Activity in the Arabidopsis Genome, Science 302 (2003) 842–846. 10.1126/science.1088305.

[20] Y. Yan, Y. Xie, Q. Gao, Y. Pan, X. Tang, Y. Liu, W. Li, H. Guo, Distinct regulation of mRNA decay pathways by ABA enhances Nitrate Reductase 1/2-derived siRNAs production and stress adaptation, Molecular Plant 18 (2025) 853–871. 10.1016/j.molp.2025.04.007.

[21] M. Zakrzewska-Placzek, F.F. Souret, G.J. Sobczyk, P.J. Green, J. Kufel, Arabidopsis thaliana XRN2 is required for primary cleavage in the pre-ribosomal RNA, Nucleic Acids Res 38 (2010) 4487–4502. 10.1093/nar/gkq172.

[22] G. Olmedo, H. Guo, B.D. Gregory, S.D. Nourizadeh, L. Aguilar-Henonin, H. Li, F. An, P. Guzman, J.R. Ecker, ETHYLENE-INSENSITIVE5 encodes a 5′→3′ exoribonuclease required for regulation of the EIN3-targeting F-box proteins EBF1/2, Proceedings of the National Academy of Sciences 103 (2006) 13286–13293. 10.1073/pnas.0605528103.

[23] G. Roman, B. Lubarsky, J.J. Kieber, M. Rothenberg, J.R. Ecker, Genetic analysis of ethylene signal transduction in Arabidopsis thaliana: five novel mutant loci integrated into a stress response pathway., Genetics 139 (1995) 1393–1409. 10.1093/genetics/139.3.1393.

[24] J.M. Alonso, A.N. Stepanova, R. Solano, E. Wisman, S. Ferrari, F.M. Ausubel, J.R. Ecker, Five components of the ethylene-response pathway identified in a screen for weak ethylene-insensitive mutants in Arabidopsis, Proceedings of the National Academy of Sciences 100 (2003) 2992–2997. 10.1073/pnas.0438070100.

[25] W. Li, M. Ma, Y. Feng, H. Li, Y. Wang, Y. Ma, M. Li, F. An, H. Guo, EIN2-Directed Translational Regulation of Ethylene Signaling in Arabidopsis, Cell 163 (2015) 670–683. 10.1016/j.cell.2015.09.037.

[26] T. Potuschak, A. Vansiri, B.M. Binder, E. Lechner, R.D. Vierstra, P. Genschik, The Exoribonuclease XRN4 Is a Component of the Ethylene Response Pathway in Arabidopsis, Plant Cell 18 (2006) 3047–3057. 10.1105/tpc.106.046508.

[27] R. Merret, V.K. Nagarajan, M.-C. Carpentier, S. Park, J.-J. Favory, J. Descombin, C. Picart, Y. Charng, P.J. Green, J.-M. Deragon, C. Bousquet-Antonelli, Heat-induced ribosome pausing triggers mRNA co-translational decay in Arabidopsis thaliana, Nucleic Acids Res 43 (2015) 4121–4132. 10.1093/nar/gkv234.

[28] T.-Q. Zhang, H. Lian, H. Tang, K. Dolezal, C.-M. Zhou, S. Yu, J.-H. Chen, Q. Chen, H. Liu, K. Ljung, J.-W. Wang, An Intrinsic MicroRNA Timer Regulates Progressive Decline in Shoot Regenerative Capacity in Plants, Plant Cell 27 (2015) 349–360. 10.1105/tpc.114.135186.

[29] F.F. Souret, J.P. Kastenmayer, P.J. Green, AtXRN4 Degrades mRNA in *Arabidopsis* and Its Substrates Include Selected miRNA Targets, Molecular Cell 15 (2004) 173–183. 10.1016/j.molcel.2004.06.006.

[30] S. Gazzani, T. Lawrenson, C. Woodward, D. Headon, R. Sablowski, A Link Between mRNA Turnover and RNA Interference in Arabidopsis, Science 306 (2004) 1046–1048. 10.1126/science.1101092.

[31] B.D. Gregory, R.C. O’Malley, R. Lister, M.A. Urich, J. Tonti-Filippini, H. Chen, A.H. Millar, J.R. Ecker, A Link between RNA Metabolism and Silencing Affecting Arabidopsis Development, Developmental Cell 14 (2008) 854–866. 10.1016/j.devcel.2008.04.005.

[32] M.A. German, M. Pillay, D.-H. Jeong, A. Hetawal, S. Luo, P. Janardhanan, V. Kannan, L.A. Rymarquis, K. Nobuta, R. German, E. De Paoli, C. Lu, G. Schroth, B.C. Meyers, P.J. Green, Global identification of microRNA–target RNA pairs by parallel analysis of RNA ends, Nat Biotechnol 26 (2008) 941–946. 10.1038/nbt1417.

[33] D.C. Amberg, A.L. Goldstein, C.N. Cole, Isolation and characterization of RAT1: an essential gene of Saccharomyces cerevisiae required for the efficient nucleocytoplasmic trafficking of mRNA., Genes Dev. 6 (1992) 1173–1189. 10.1101/gad.6.7.1173.

[34] F. He, X. Li, P. Spatrick, R. Casillo, S. Dong, A. Jacobson, Genome-Wide Analysis of mRNAs Regulated by the Nonsense-Mediated and 5′ to 3′ mRNA Decay Pathways in Yeast, Molecular Cell 12 (2003) 1439–1452. 10.1016/S1097-2765(03)00446-5.

[35] M. Jinek, S.M. Coyle, J.A. Doudna, Coupled 5′ Nucleotide Recognition and Processivity in Xrn1-Mediated mRNA Decay, Molecular Cell 41 (2011) 600–608. 10.1016/j.molcel.2011.02.004.

[36] C.I. Jones, M.V. Zabolotskaya, S.F. Newbury, The 5′ → 3′ exoribonuclease XRN1/Pacman and its functions in cellular processes and development, WIREs RNA 3 (2012) 455–468. 10.1002/wrna.1109.

[37] M. Kenna, A. Stevens, M. McCammon, M.G. Douglas, An essential yeast gene with homology to the exonuclease-encoding XRN1/KEM1 gene also encodes a protein with exoribonuclease activity., Mol Cell Biol 13 (1993) 341–350. 10.1128/mcb.13.1.341.

[38] A.M. Page, K. Davis, C. Molineux, R.D. Kolodner, A.W. Johnson, Mutational analysis of exoribonuclease I from Saccharomyces cerevisiae, Nucleic Acids Res 26 (1998) 3707–3716. 10.1093/nar/26.16.3707.

[39] L.-H. Qu, A. Henras, Y.-J. Lu, H. Zhou, W. Zhou, Y.-Q. Zhu, J. Zhao, Y. Henry, M. Caizergues-Ferrer, J.-P. Bachellerie, Seven Novel Methylation Guide Small Nucleolar RNAs Are Processed from a Common Polycistronic Transcript by Rat1p and RNase III in Yeast, Mol Cell Biol 19 (1999) 1144–1158. 10.1128/mcb.19.2.1144.

[40] J.A. Solinger, D. Pascolini, W.-D. Heyer, Active-Site Mutations in the Xrn1p Exoribonuclease of Saccharomyces cerevisiae Reveal a Specific Role in Meiosis, Molecular and Cellular Biology 19 (1999) 5930–5942. 10.1128/MCB.19.9.5930.

[41] P. Tesina, E. Heckel, J. Cheng, M. Fromont-Racine, R. Buschauer, L. Kater, B. Beatrix, O. Berninghausen, A. Jacquier, T. Becker, R. Beckmann, Structure of the 80S ribosome–Xrn1 nuclease complex, Nat Struct Mol Biol 26 (2019) 275–280. 10.1038/s41594-019-0202-5.

[42] Y. Zeng, H.-W. Zhang, X.-X. Wu, Y. Zhang, Structural basis of exoribonuclease-mediated mRNA transcription termination, Nature 628 (2024) 887–893. 10.1038/s41586-024-07240-3.

[43] M.T. Panteva, G.M. Giambaşu, D.M. York, Comparison of structural, thermodynamic, kinetic and mass transport properties of Mg2+ ion models commonly used in biomolecular simulations, Journal of Computational Chemistry 36 (2015) 970–982. 10.1002/jcc.23881.

[44] J.G. Gezelle, S.M. Korn, J.T. McDonald, Z. Gong, A. Erickson, C.-H. Huang, F. Yang, M. Cronin, Y.-W. Kuo, B.T. Wimberly, A.-L. Steckelberg, A conserved viral RNA fold enables nuclease resistance across kingdoms of life, Nucleic Acids Res 53 (2025) gkaf840. 10.1093/nar/gkaf840.

[45] M.J. Szucs, P.J. Nichols, R.A. Jones, Q. Vicens, J.S. Kieft, A New Subclass of Exoribonuclease-Resistant RNA Found in Multiple Genera of Flaviviridae, mBio 11 (2020) 10.1128/mbio.02352-20. 10.1128/mbio.02352-20.

[46] Q. Vicens, J.S. Kieft, Shared properties and singularities of exoribonuclease-resistant RNAs in viruses, Computational and Structural Biotechnology Journal 19 (2021) 4373–4380. 10.1016/j.csbj.2021.07.024.

[47] J.H. Chang, S. Xiang, K. Xiang, J.L. Manley, L. Tong, Structural and biochemical studies of the 5′→3′ exoribonuclease Xrn1, Nat Struct Mol Biol 18 (2011) 270–276. 10.1038/nsmb.1984.

[48] S.F. Altschul, T.L. Madden, A.A. Schäffer, J. Zhang, Z. Zhang, W. Miller, D.J. Lipman, Gapped BLAST and PSI-BLAST: a new generation of protein database search programs, Nucleic Acids Res 25 (1997) 3389–3402. 10.1093/nar/25.17.3389.

[49] A.A. Schäffer, L. Aravind, T.L. Madden, S. Shavirin, J.L. Spouge, Y.I. Wolf, E.V. Koonin, S.F. Altschul, Improving the accuracy of PSI-BLAST protein database searches with composition-based statistics and other refinements, Nucleic Acids Res 29 (2001) 2994–3005. 10.1093/nar/29.14.2994.

[50] The UniProt Consortium, UniProt: the Universal Protein Knowledgebase in 2025, Nucleic Acids Res 53 (2025) D609–D617. 10.1093/nar/gkae1010.

[51] UniProt, (2026). https://www.uniprot.org (accessed May 6, 2026).

[52] F. Lemoine, D. Correia, V. Lefort, O. Doppelt-Azeroual, F. Mareuil, S. Cohen-Boulakia, O. Gascuel, NGPhylogeny.fr: new generation phylogenetic services for non-specialists, Nucleic Acids Res 47 (2019) W260–W265. 10.1093/nar/gkz303.

[53] NGPhylogeny.fr, (2026). https://ngphylogeny.fr/ (accessed July 10, 2026).

[54] F. Schulz, J. Andreani, R. Francis, H. Boudjemaa, J.Y. Bou Khalil, J. Lee, B. La Scola, T. Woyke, Advantages and Limits of Metagenomic Assembly and Binning of a Giant Virus, mSystems 5 (2020) 10.1128/msystems.00048-20. 10.1128/msystems.00048-20.

[55] F. Sievers, A. Wilm, D. Dineen, T.J. Gibson, K. Karplus, W. Li, R. Lopez, H. McWilliam, M. Remmert, J. Söding, J.D. Thompson, D.G. Higgins, Fast, scalable generation of high-quality protein multiple sequence alignments using Clustal Omega, Molecular Systems Biology (2011). 10.1038/msb.2011.75.

[56] A. Criscuolo, S. Gribaldo, BMGE (Block Mapping and Gathering with Entropy): a new software for selection of phylogenetic informative regions from multiple sequence alignments, BMC Evol Biol 10 (2010) 210. 10.1186/1471-2148-10-210.

[57] S. Guindon, J.-F. Dufayard, V. Lefort, M. Anisimova, W. Hordijk, O. Gascuel, New Algorithms and Methods to Estimate Maximum-Likelihood Phylogenies: Assessing the Performance of PhyML 3.0, Syst Biol 59 (2010) 307–321. 10.1093/sysbio/syq010.

[58] V. Lefort, J.-E. Longueville, O. Gascuel, SMS: Smart Model Selection in PhyML, Mol Biol Evol 34 (2017) 2422–2424. 10.1093/molbev/msx149.

[59] T. Junier, E.M. Zdobnov, The Newick utilities: high-throughput phylogenetic tree processing in the Unix shell, Bioinformatics 26 (2010) 1669–1670. 10.1093/bioinformatics/btq243.

[60] I. Letunic, P. Bork, Interactive Tree of Life (iTOL) v6: recent updates to the phylogenetic tree display and annotation tool, Nucleic Acids Res 52 (2024) W78–W82. 10.1093/nar/gkae268.

[61] iTOL, (2026). https://itol.embl.de/ (accessed July 10, 2026).

[62] C. Discovery, J. Boitreaud, J. Dent, M. McPartlon, J. Meier, V. Reis, A. Rogozhnikov, K. Wu, Chai-1: Decoding the molecular interactions of life, (2024) 2024.10.10.615955. 10.1101/2024.10.10.615955.

[63] S. Passaro, G. Corso, J. Wohlwend, M. Reveiz, S. Thaler, V.R. Somnath, N. Getz, T. Portnoi, J. Roy, H. Stark, D. Kwabi-Addo, D. Beaini, T. Jaakkola, R. Barzilay, Boltz-2: Towards Accurate and Efficient Binding Affinity Prediction, (2025) 2025.06.14.659707. 10.1101/2025.06.14.659707.

[64] Tamarind Bio, Tamarind Bio (2026). https://app.tamarind.bio/app (accessed July 19, 2026).

[65] AlphaFold3, (2026). https://alphafoldserver.com/ (accessed July 15, 2026).

[66] Schrödinger, LLC, The PyMOL Molecular Graphics System, Version 3.1.5.1, (2015).

[67] U. Höweler, MAXIMOBY/MOBY, version 25.02 (12/25). CHEOPS Molecular Modelling, (2026). http://www.cheops-molecular-modeling.de.

[68] S.J. Weiner, P.A. Kollman, D.A. Case, U.C. Singh, C. Ghio, G. Alagona, S. Profeta, P. Weiner, A new force field for molecular mechanical simulation of nucleic acids and proteins, Journal American Chemical Society 106 (1984) 765–784. 10.1021/ja00315a051.

[69] J.E. Nielsen, G. Vriend, Optimizing the hydrogen-bond network in Poisson–Boltzmann equation-based pKa calculations, Proteins: Structure, Function, and Bioinformatics 43 (2001) 403–412. 10.1002/prot.1053.

[70] M. Scherlo, E. Wippermann, T. Fürtges, R. Künne, A. Yelboğa, F. Rütten, M. Böckmann, U. Höweler, T. Rudack, kontakteUR: transforming coordinates to chemical intuition to focus on essential interactions in biomolecular systems, (2026) 2026.06.19.732925. 10.64898/2026.06.19.732925.

[71] T. Williams, C. Kelley, Gnuplot, version 5.4, (2020). https://gnuplot.sourceforge.net/.

[72] M.C. Frith, N.F.W. Saunders, B. Kobe, T.L. Bailey, Discovering Sequence Motifs with Arbitrary Insertions and Deletions, PLOS Computational Biology 4 (2008) e1000071. 10.1371/journal.pcbi.1000071.

[73] MEME Suite, (2026). https://meme-suite.org/meme/index.html (accessed July 10, 2026).

[74] F. Madeira, N. Madhusoodanan, J. Lee, A. Eusebi, A. Niewielska, A.R.N. Tivey, R. Lopez, S. Butcher, The EMBL-EBI Job Dispatcher sequence analysis tools framework in 2024, Nucleic Acids Res 52 (2024) W521–W525. 10.1093/nar/gkae241.

[75] web server EMBL-EBI, E.B. Institute, Clustal Omega, (2026). https://www.ebi.ac.uk/jdispatcher/msa/clustalo (accessed July 10, 2026).

[76] H. Nielsen, F. Teufel, S. Brunak, G. von Heijne, SignalP: The Evolution of a Web Server, in: F. Lisacek (Ed.), Protein Bioinformatics, Springer US, New York, NY, 2024: pp. 331–367. 10.1007/978-1-0716-4007-4_17.

[77] web server DTU Health Tech, SignalP-6.0, (2026). https://services.healthtech.dtu.dk/services/SignalP-6.0/ (accessed July 10, 2026).

[78] A. Kryshtafovych, T. Schwede, M. Topf, K. Fidelis, J. Moult, Progress and Bottlenecks for Deep Learning in Computational Structure Biology: CASP Round XVI, Proteins: Structure, Function, and Bioinformatics 94 (2026) 5–14. 10.1002/prot.70076.

[79] A. Kryshtafovych, T. Schwede, M. Topf, K. Fidelis, J. Moult, Critical assessment of methods of protein structure prediction (CASP)—Round XV, Proteins: Structure, Function, and Bioinformatics 91 (2023) 1539–1549. 10.1002/prot.26617.

[80] A. Kryshtafovych, T. Schwede, M. Topf, K. Fidelis, J. Moult, Critical assessment of methods of protein structure prediction (CASP)—Round XIV, Proteins: Structure, Function, and Bioinformatics 89 (2021) 1607–1617. 10.1002/prot.26237.

[81] A. Kryshtafovych, T. Schwede, M. Topf, K. Fidelis, J. Moult, Critical assessment of methods of protein structure prediction (CASP)—Round XIII, Proteins: Structure, Function, and Bioinformatics 87 (2019) 1011–1020. 10.1002/prot.25823.

[82] A.W. Senior, R. Evans, J. Jumper, J. Kirkpatrick, L. Sifre, T. Green, C. Qin, A. Žídek, A.W.R. Nelson, A. Bridgland, H. Penedones, S. Petersen, K. Simonyan, S. Crossan, P. Kohli, D.T. Jones, D. Silver, K. Kavukcuoglu, D. Hassabis, Improved protein structure prediction using potentials from deep learning, Nature 577 (2020) 706–710. 10.1038/s41586-019-1923-7.

[83] R.C. Kretsch, A.M. Hummer, S. He, R. Yuan, J. Zhang, T. Karagianes, Q. Cong, A. Kryshtafovych, R. Das, Assessment of Nucleic Acid Structure Prediction in CASP16, Proteins: Structure, Function, and Bioinformatics n/a (2025). 10.1002/prot.70072.

[84] M. Kim, N.J. Krogan, L. Vasiljeva, O.J. Rando, E. Nedea, J.F. Greenblatt, S. Buratowski, The yeast Rat1 exonuclease promotes transcription termination by RNA polymerase II, Nature 432 (2004) 517–522. 10.1038/nature03041.

[85] J.H. Overbeck, D. Stelzig, A.-L. Fuchs, J.P. Wurm, R. Sprangers, Observation of conformational changes that underlie the catalytic cycle of Xrn2, Nat Chem Biol 18 (2022) 1152–1160. 10.1038/s41589-022-01111-6.

[86] G.C. MacIntosh, M.S. Hillwig, A. Meyer, L. Flagel, RNase T2 genes from rice and the evolution of secretory ribonucleases in plants, Mol Genet Genomics 283 (2010) 381–396. 10.1007/s00438-010-0524-9.

[87] F. Cheng, E. Zamski, W. Guo, D.M. Pharr, J.D. Williamson, Salicylic acid stimulates secretion of the normally symplastic enzyme mannitol dehydrogenase: a possible defense against mannitol-secreting fungal pathogens, Planta 230 (2009) 1093–1103. 10.1007/s00425-009-1006-3.

[88] Y. Ding, J. Wang, J. Wang, Y.-D. Stierhof, D.G. Robinson, L. Jiang, Unconventional protein secretion, Trends in Plant Science 17 (2012) 606–615. 10.1016/j.tplants.2012.06.004.

