## Supplemental information for "Computational and structure-based analysis of the conserved 5′–3′ exoribonuclease (XRN) family in plant species suggests seven potentially functional amino acid motifs within the N-terminal catalytic core"

**Supplemental figures**

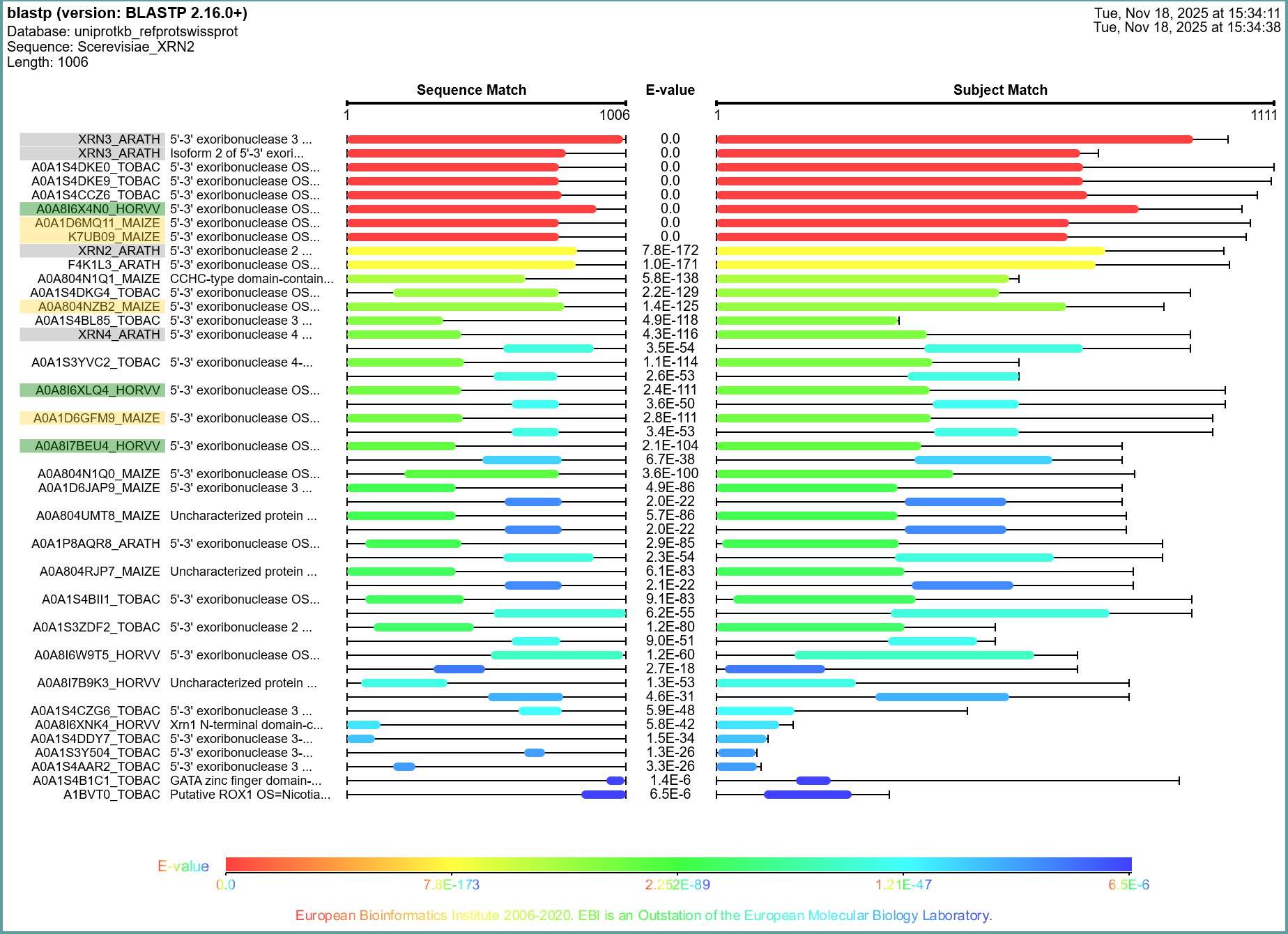

Figure S 1. XRN candidates identified by BLASTP.

Names of candidate sequences that were selected for further analysis are shaded in different colors: Arabidopsis enzymes in grey, protein sequences from maize in yellow, and barley in green. Results are sorted from lowest to highest E-value.

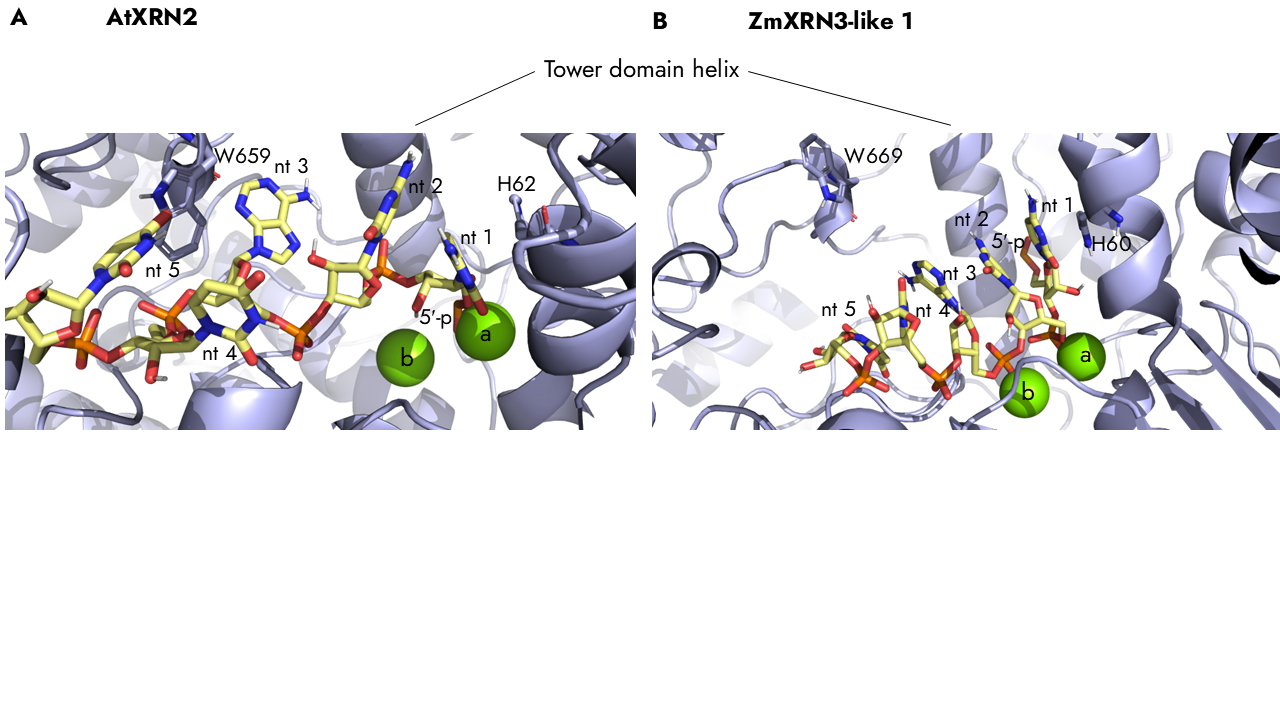

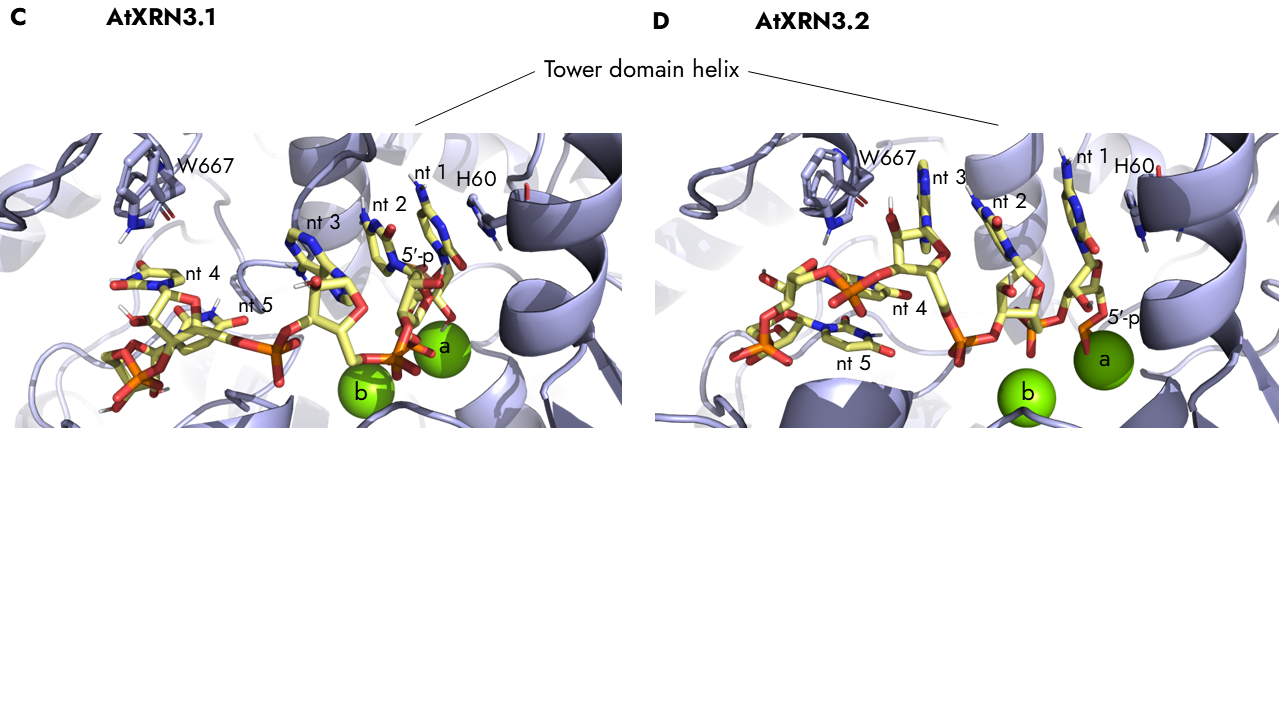

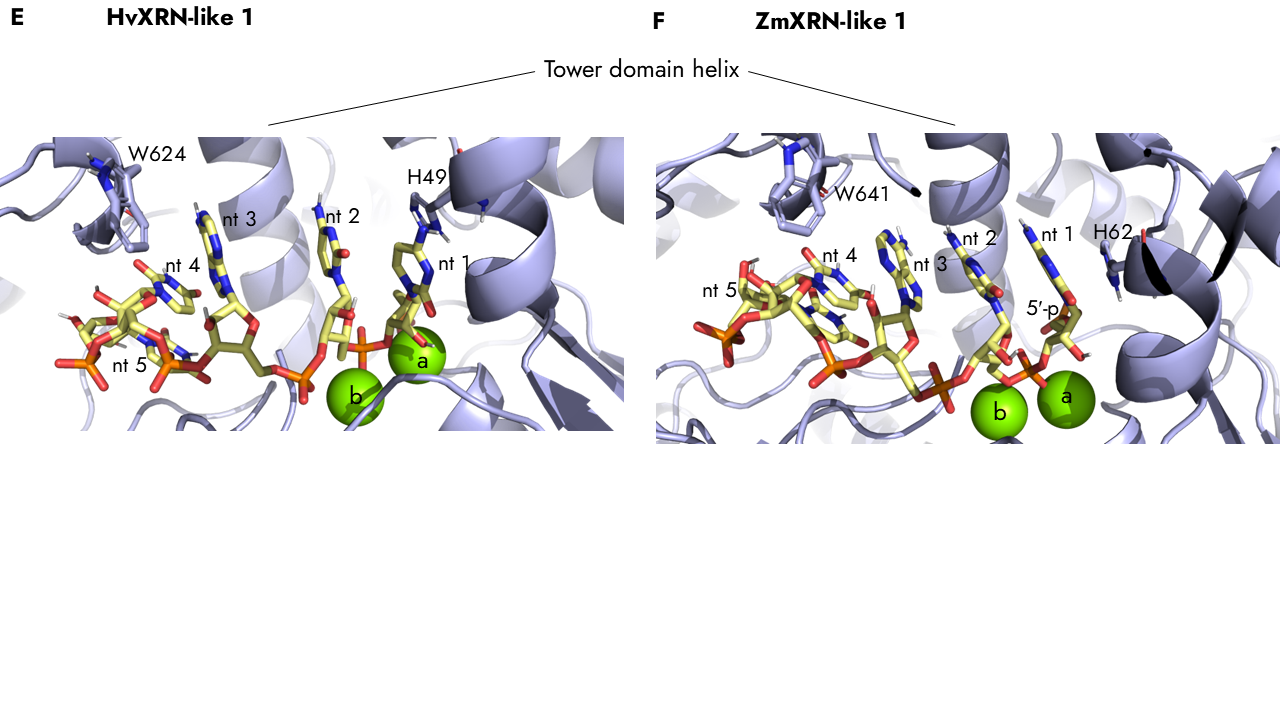

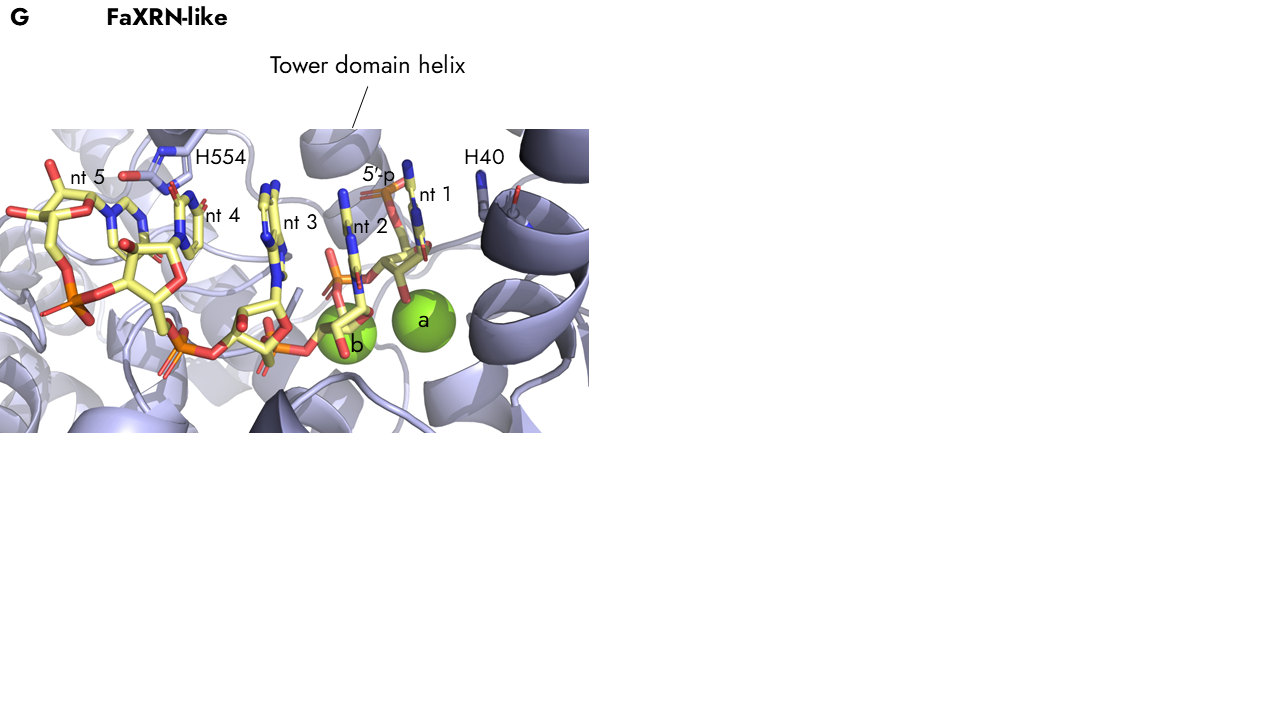

Figure S 2. Conserved active center of the XRN2/3 clade, the monocot XRN clade, and FaXRN-like predicted by structure prediction. The protein structures were predicted with a 5 nt long ssRNA and two magnesium ions (a, b) by Chai-1 and further processed via MAXIMOBY. An overview of the enzyme structure is shown. Enzyme in light blue, ssRNA in yellow, atoms colored by element: hydrogen in white, nitrogen in blue, oxygen in red, sulfur in orange, and magnesium in green. W: tryptophan, H: histidine, 5′-p: 5′-monophosphate, nt: nucleotide. A. AtXRN2, B. ZmXRN3-like 1, C. AtXRN3.1, D. AtXRN3.2, E. HvXRN-like 1, F. ZmXRN-like 1, G. FaXRN-like.

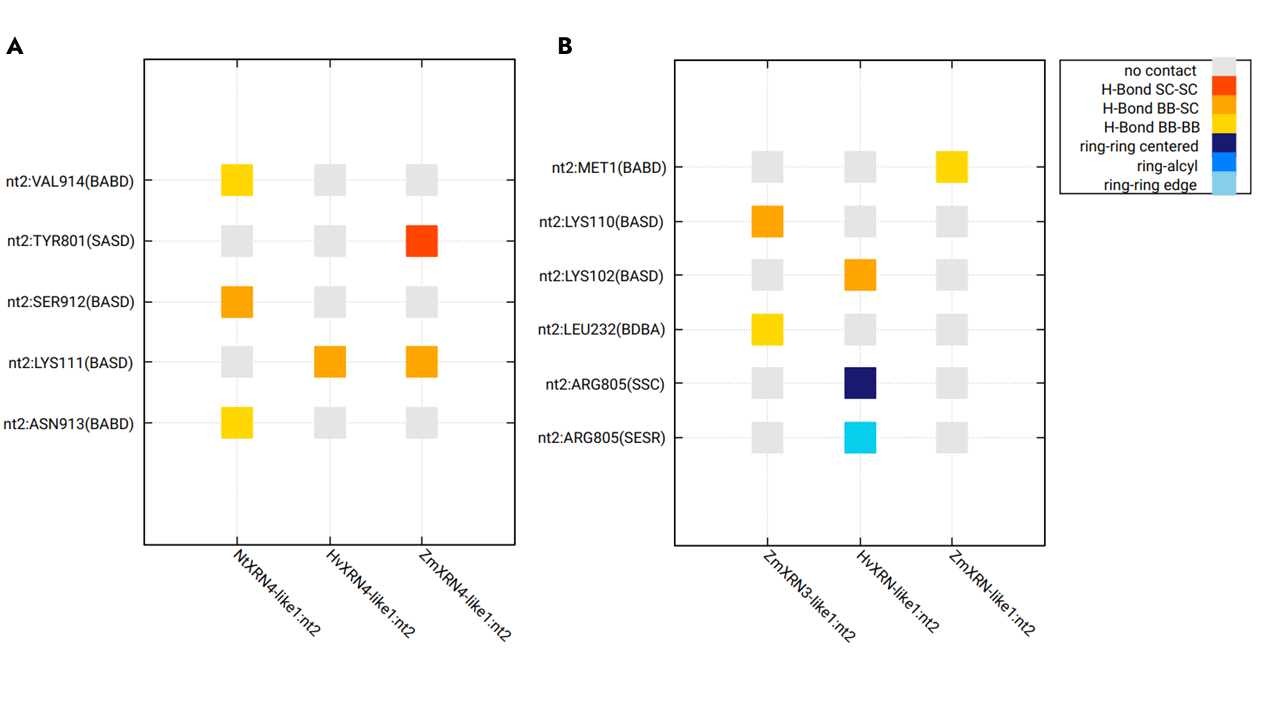

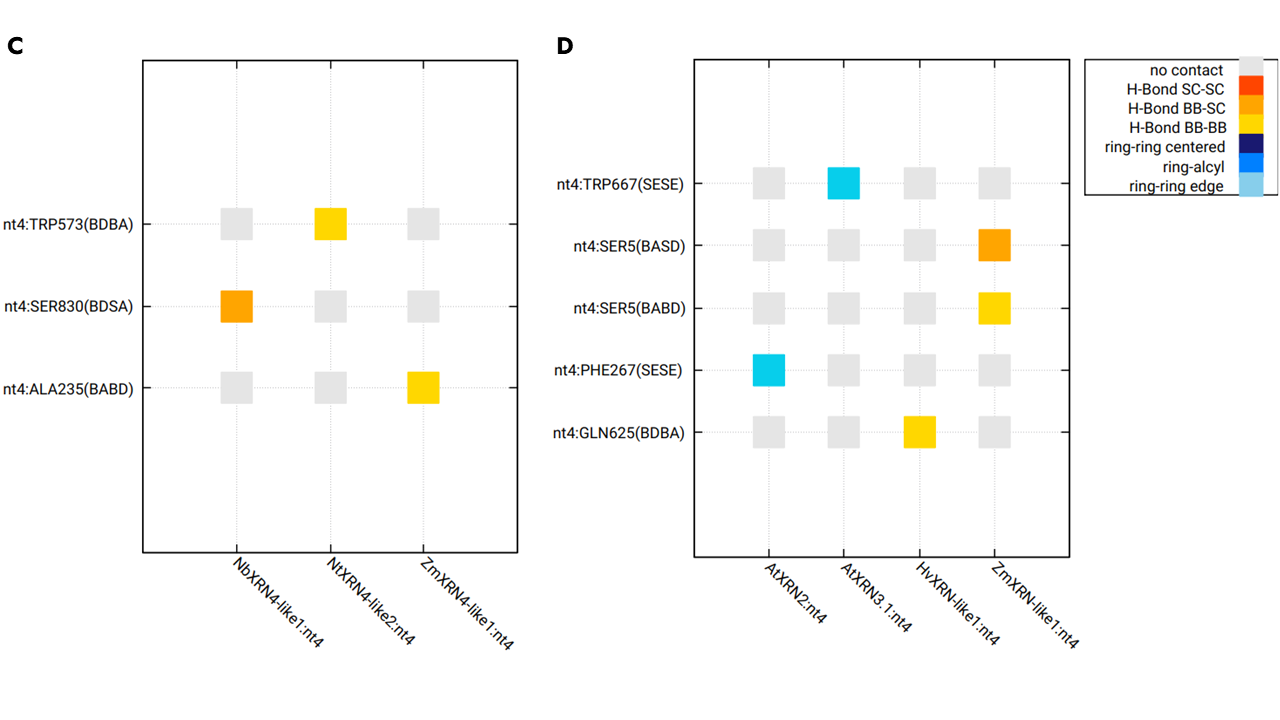

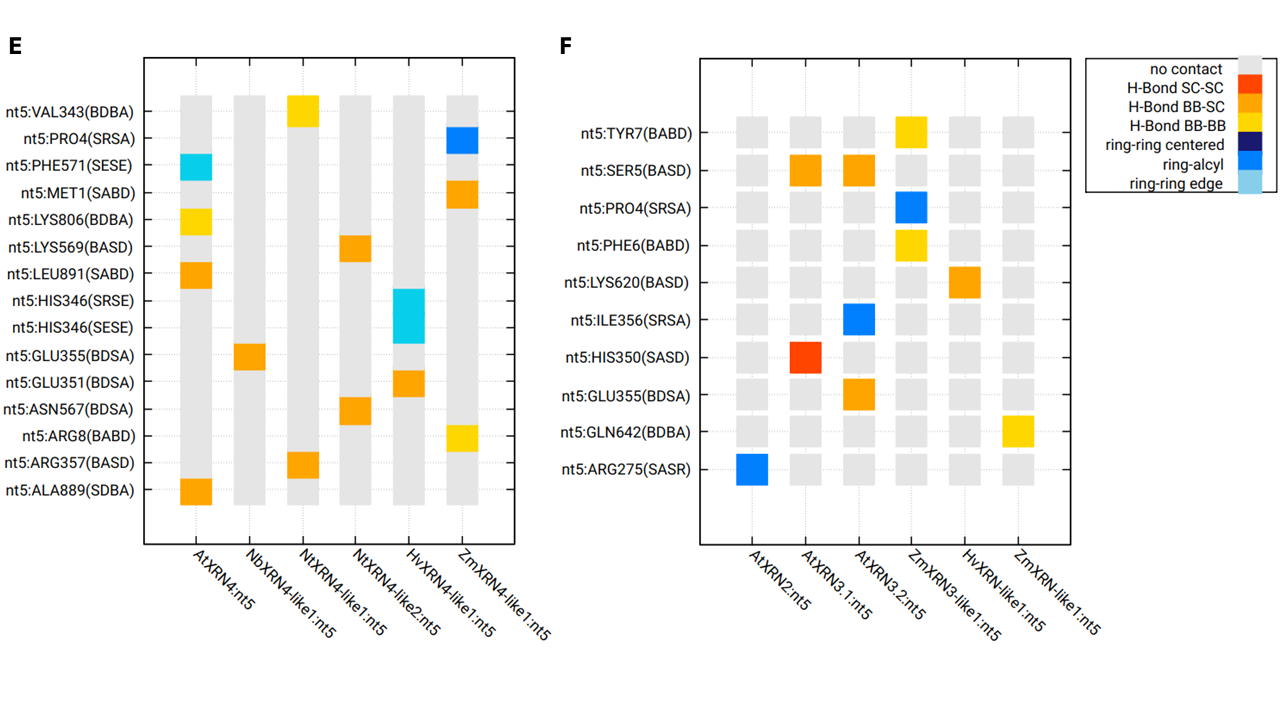

Figure S 3. Contact analysis revealed amino acids for ssRNA mediation. A, B. Contacts of nt 2 with A. the XRN4-like clade models and B. the models of the XRN2/3-like clade and the monocot XRN-like clade. C, D. Contacts of nt 4 with C. the XRN4-like clade models and D. the models of the XRN2/3-like clade and the monocot XRN-like clade. E, F. Contacts of nt 5 with E. the XRN4-like clade models and F. the models of the XRN2/3-like clade and the monocot XRN-like clade. SC in legend: side chain, BB in legend: backbone, Abbreviations of the y-axis: S: side chain, B: backbone. In hydrogen bonds: H: hydrogen, D: H-donor, A: H-acceptor. In ring interactions: R: ring (face), E: ring (edge), A: alkyl group, C: ring (center). nt: nucleotide.

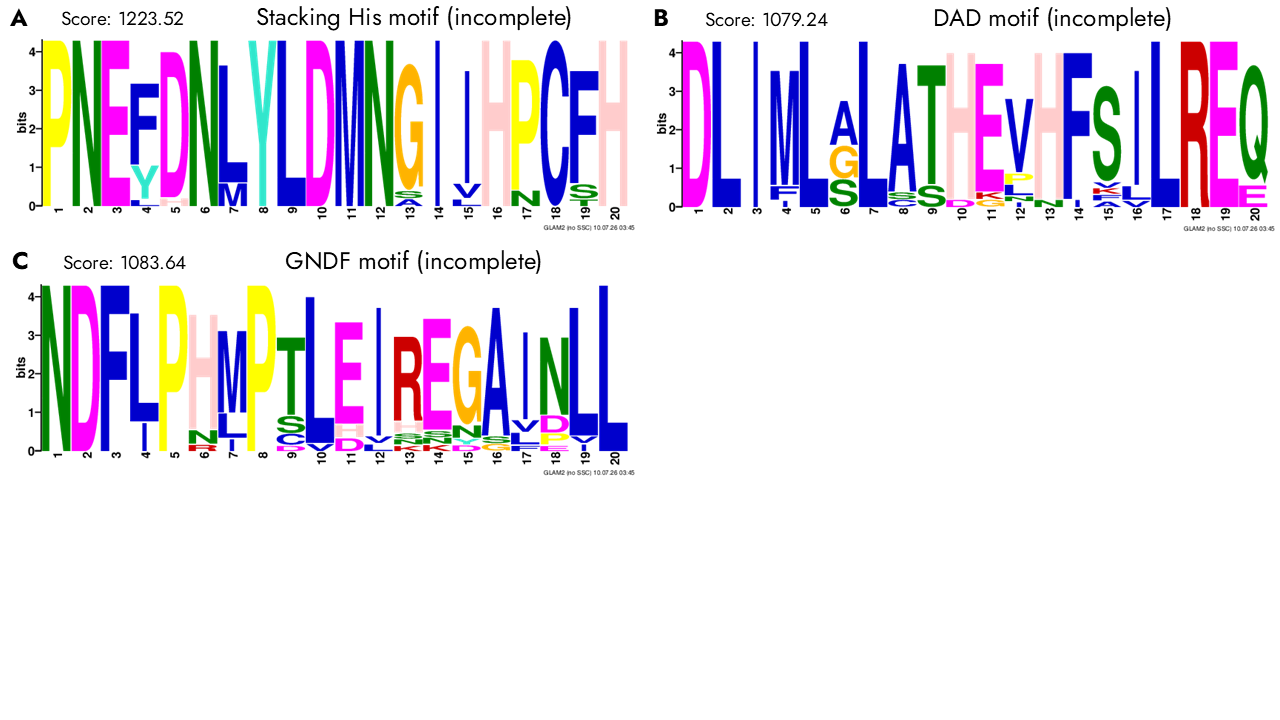

Figure S 4. Sequence motif screening showed three incomplete motifs. **A**. Stacking His motif. **B**. DAD motif. **C**. GNDF motif.

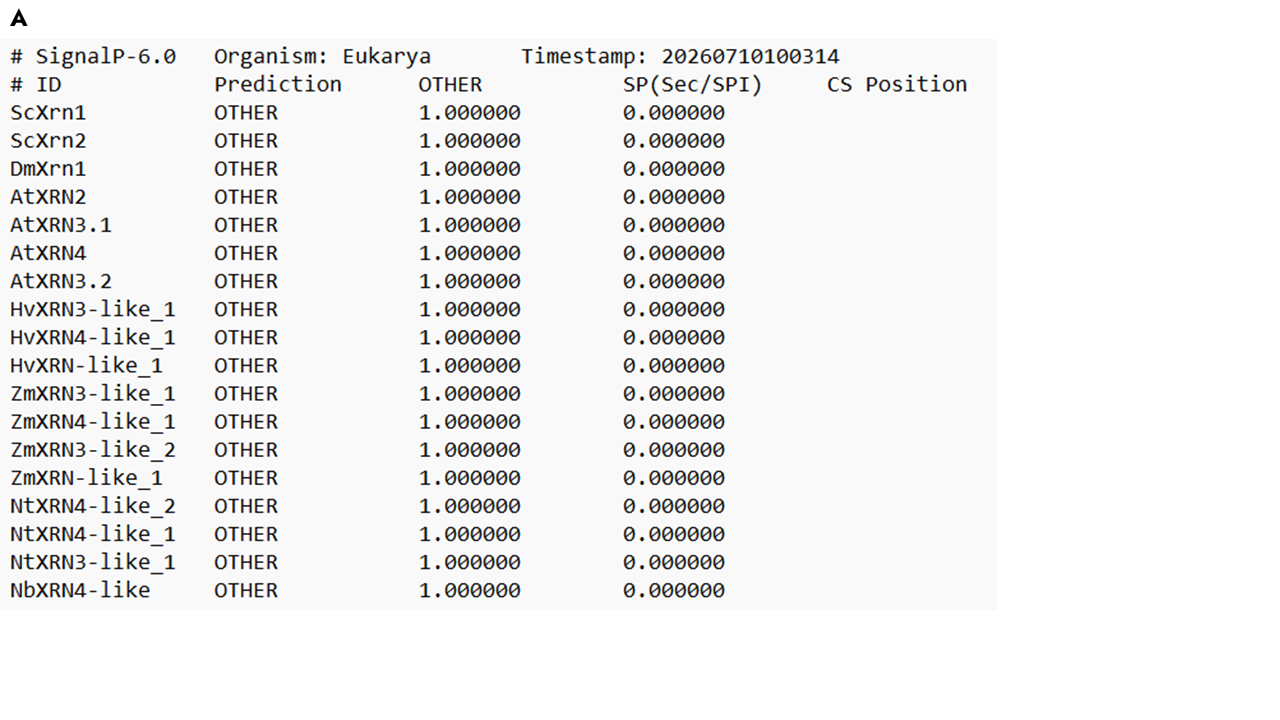

Figure S 5. Sequence motif analysis suggests no transport of the plant XRNs into the apoplast. SignalP prediction shows no secretory signal peptides (SP) for the investigated XRN sequences. SP (Sec/SPI): secretory signal peptides transported by the Sec translocon and cleaved by Signal Peptidase I (*Lep*).

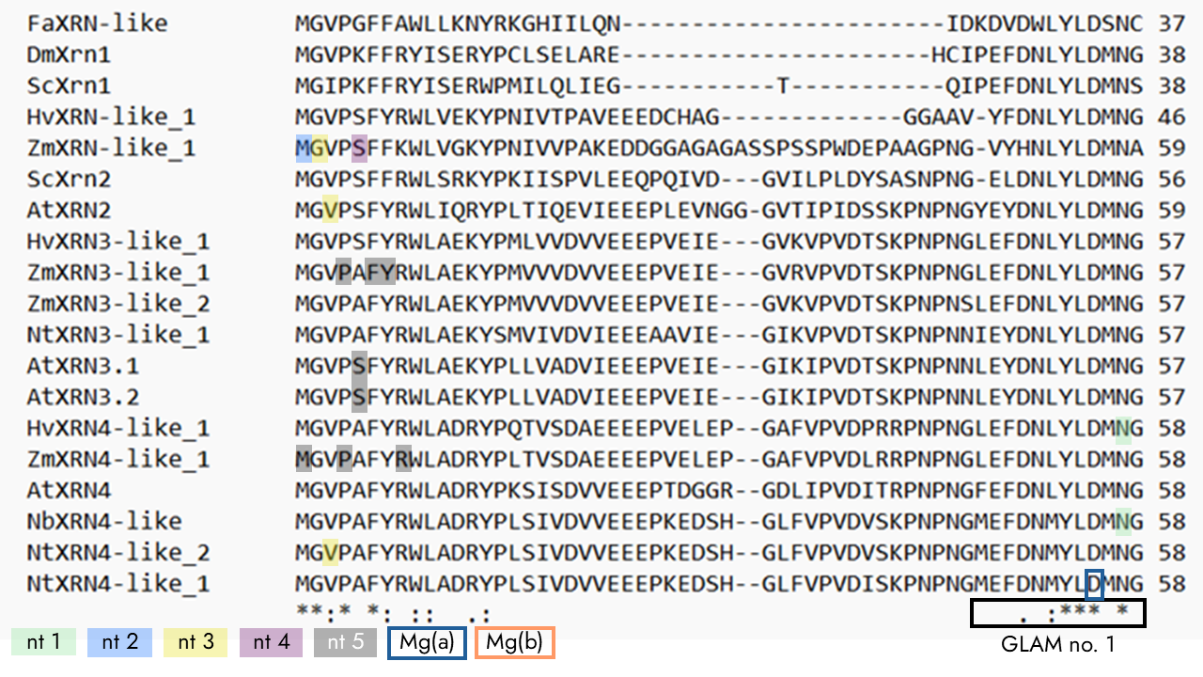

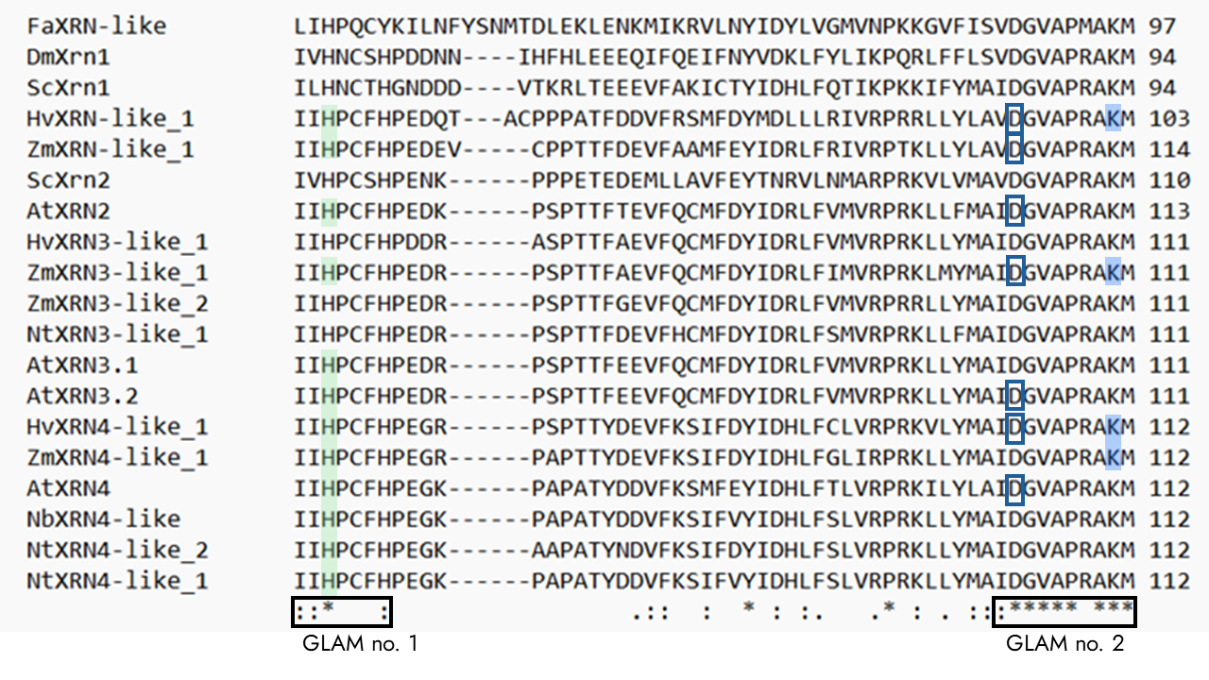

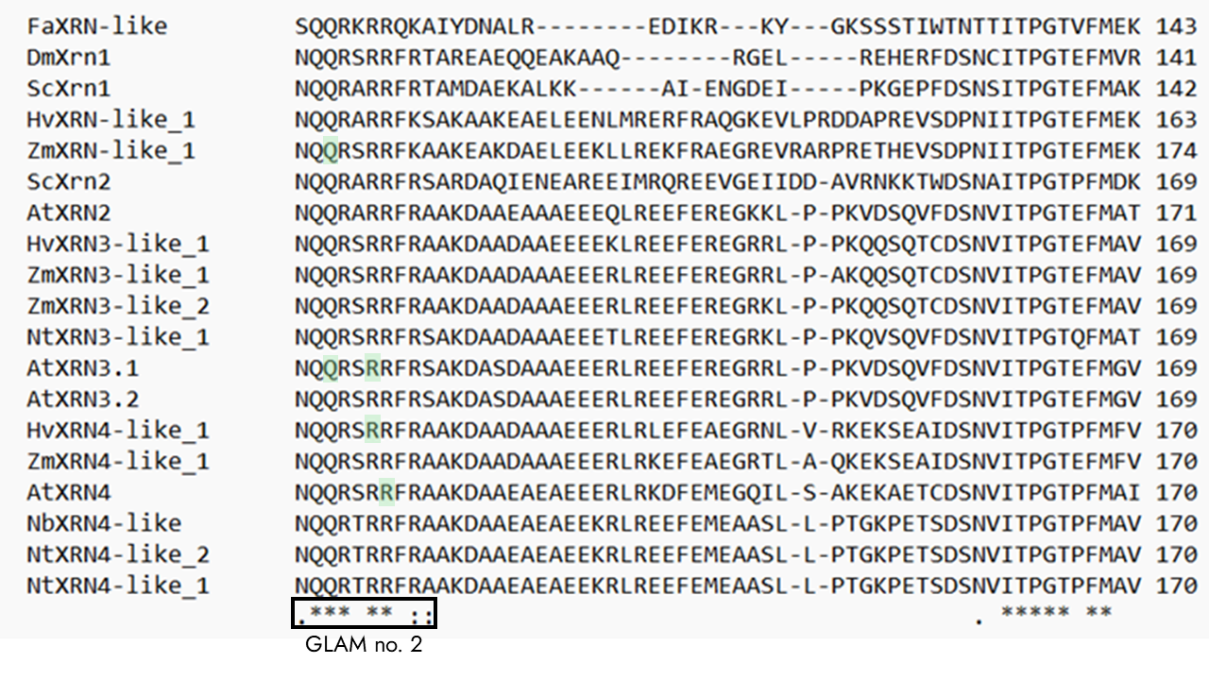

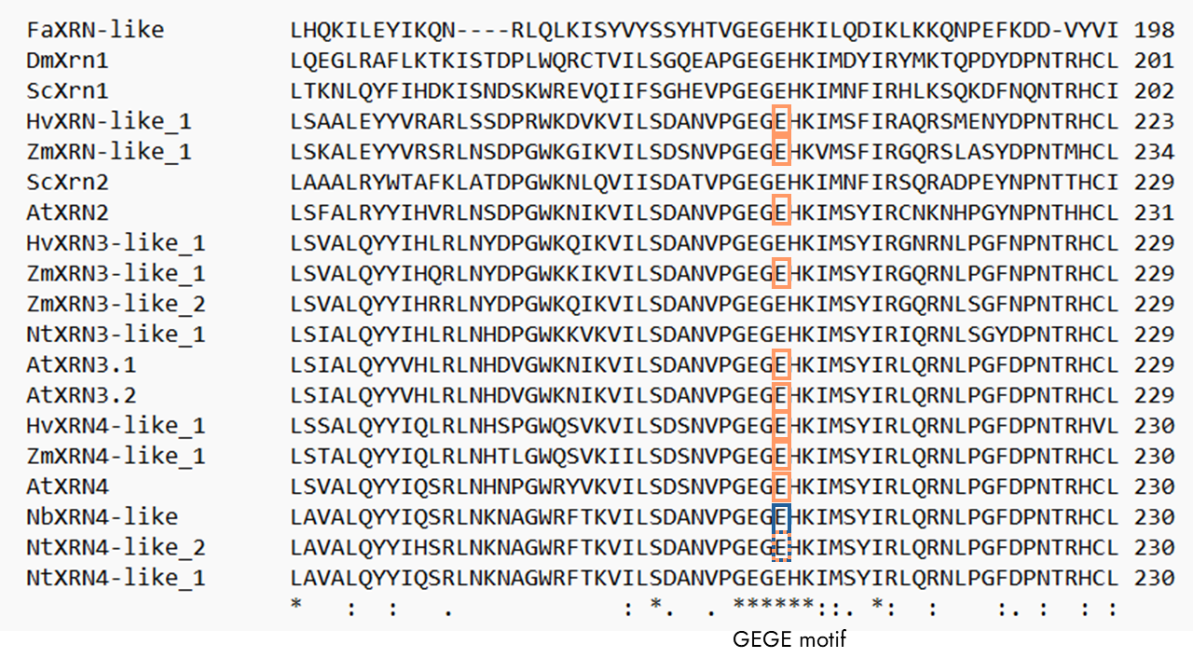

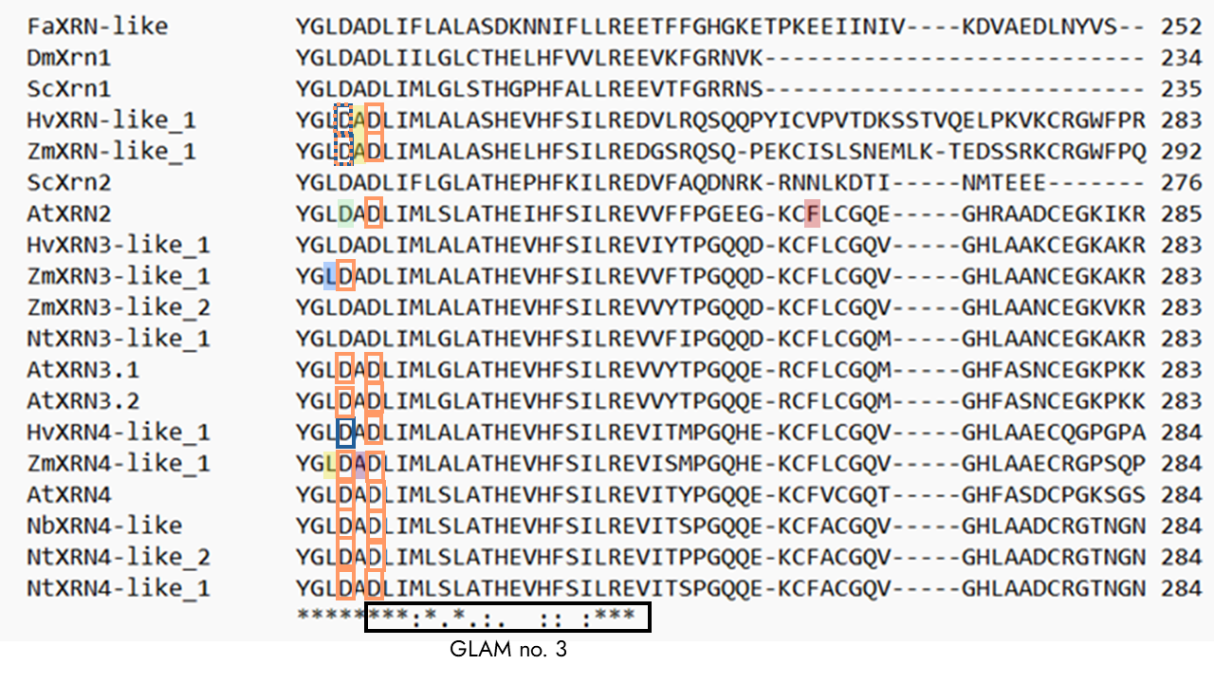

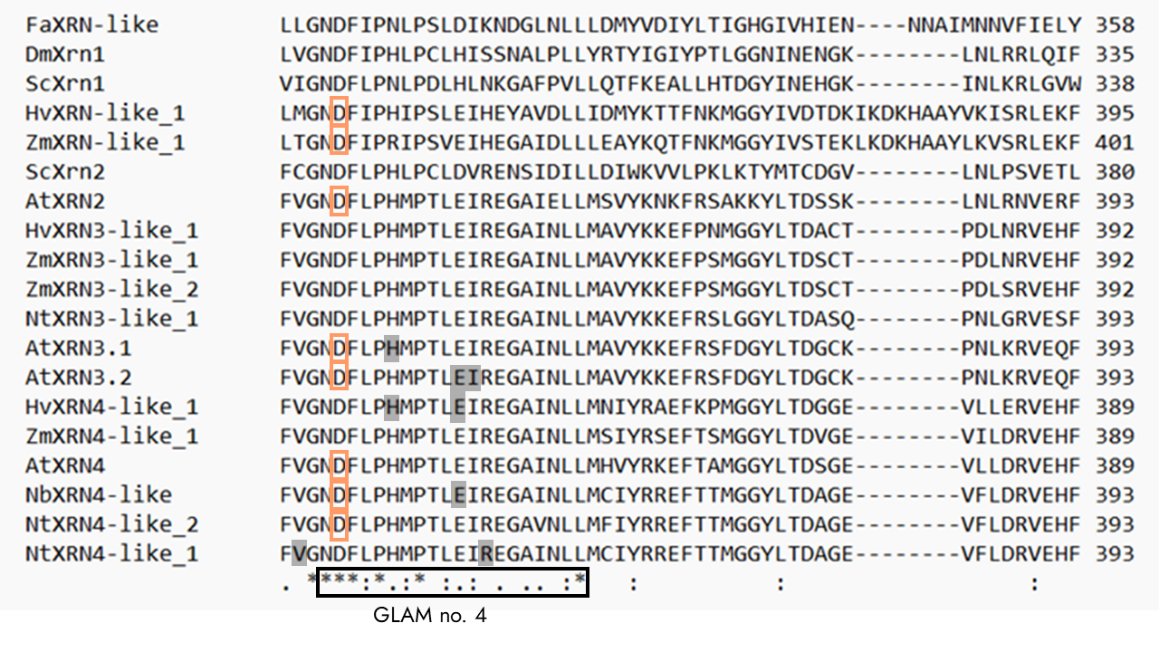

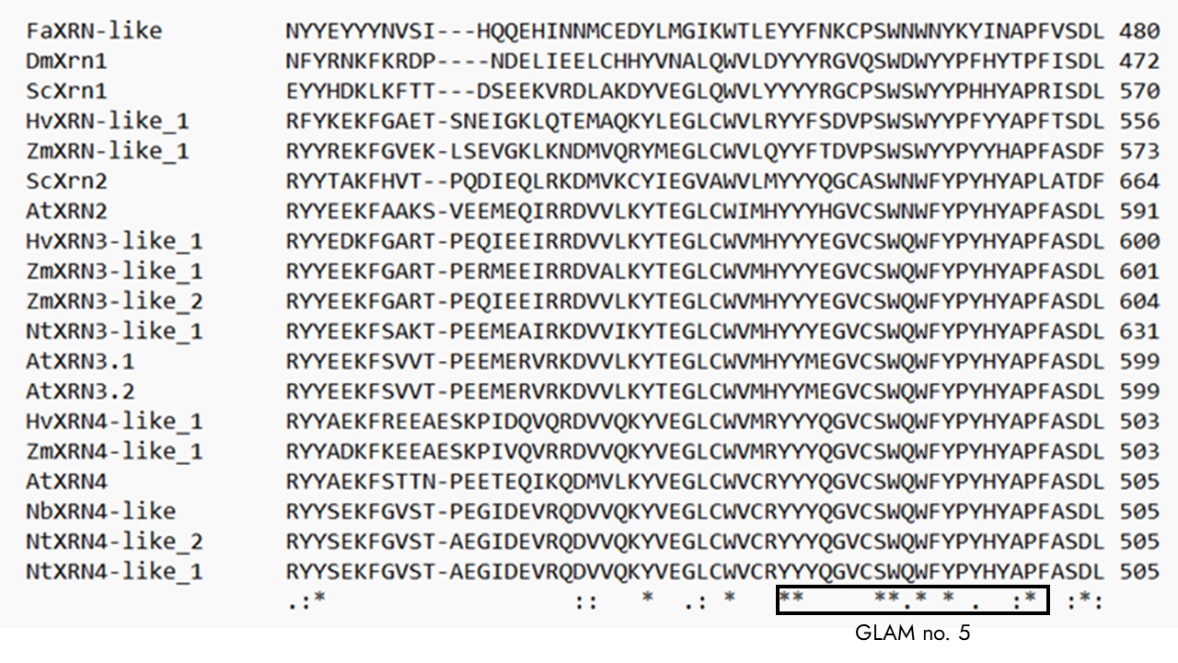

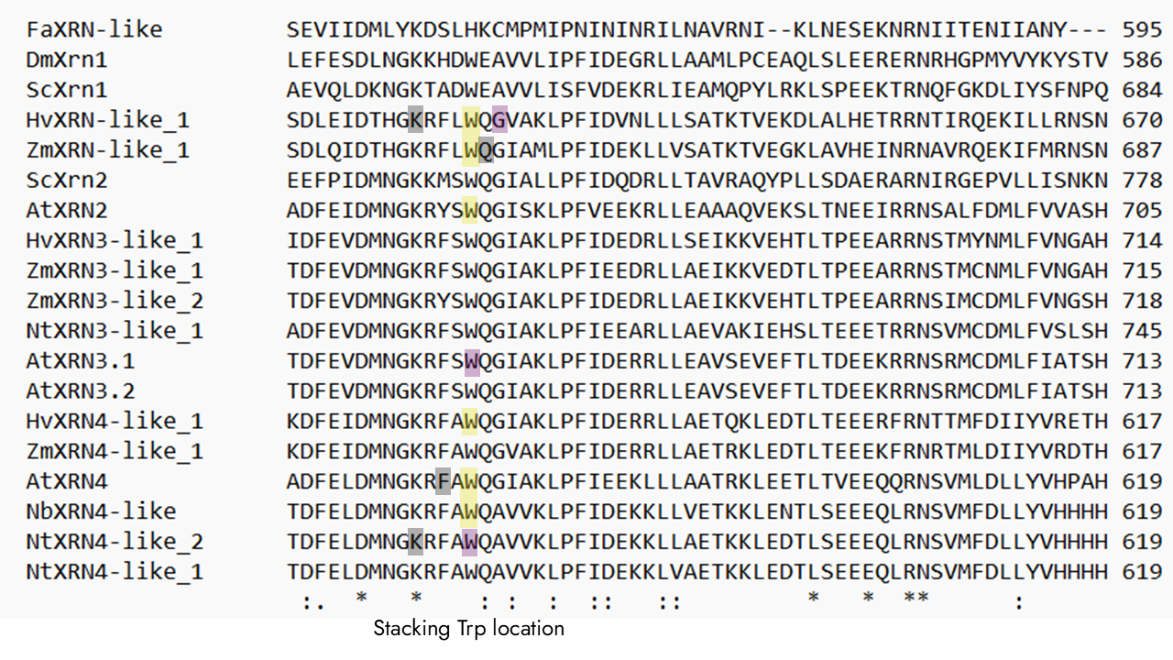

Figure S 6. MSA with amino acids from contact analysis. GLAM motif numbers (no.) marked. Conserved amino acids are indicated with an asterisk.

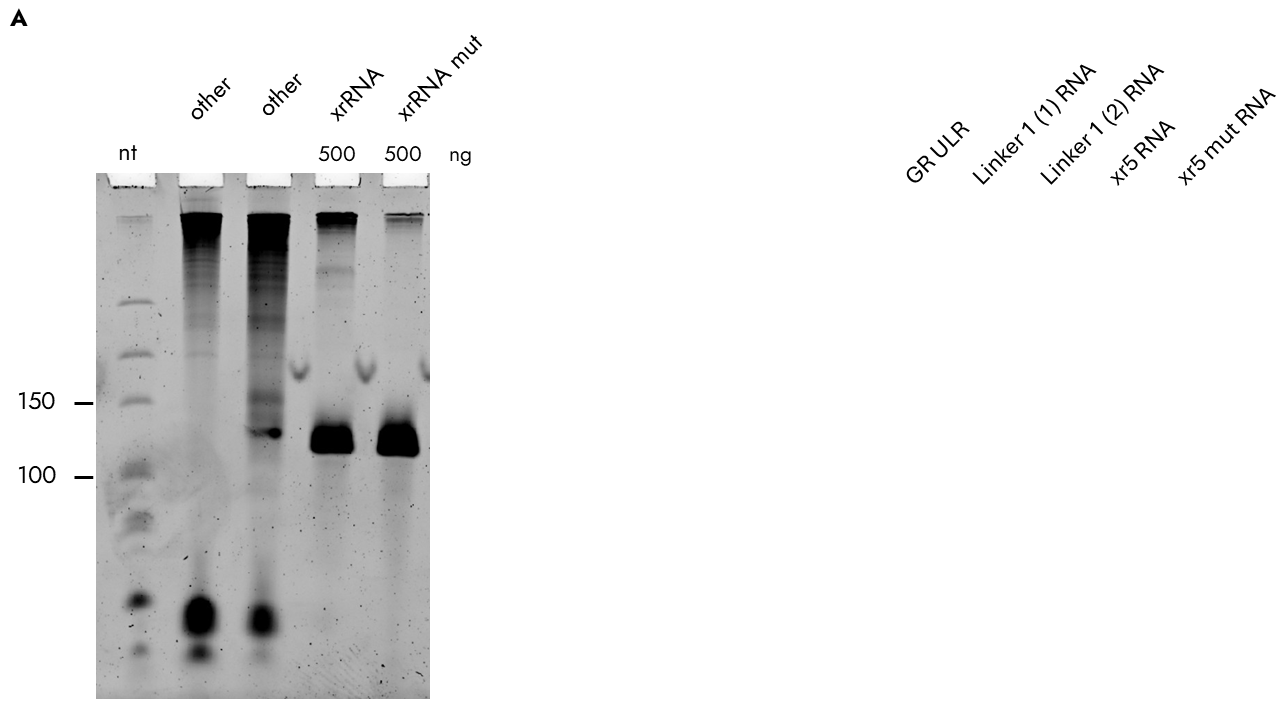

Figure S 7. 6% urea polyacrylamide gel showing the xrRNA design without and with two point mutations in the L2A region at its expected size of 114 nt. mut: mutated, nt: nucleotide, ng: nanograms, other: irrelevant samples.

**Supplemental tables**

Table S 1. Accession numbers of all protein sequences used.

| **Phylogenetic Designation** | **UniProt Accession** | **Organisms** | **Gene-ID/ Accession** | **Assembly** | **Protein length in [aa]** |
| --- | --- | --- | --- | --- | --- |
| ScXRN1 | P22147 | *S. cerevisiae* | 852702 | GCF_000146045.2 | 1,528 |
| ScXRN2 | Q02792 | *S. cerevisiae* | 854213 | GCF_000146045.2 | 1,006 |
| DmXRN1 | E1JJR3 | *D. melanogaster* | 32935 | GCF_000001215.4 | 1,613 |
| AtXRN4 | Q9FQ04 | *A. thaliana* | 841891 | GCF_000001735.4 | 947 |
| NbXRN4-like 1 | A0A3S6GZK5 | *N. benthamiana* | Nbe.v1.1.chr10g09300 / KY402212 | Nbe_v1.1 | 982 |
| NtXRN4-like 1 | A0AC58S8N5 | *N. tabacum* | 107808663 | GCF_000715075.1 | 982 |
| NtXRN4-like 2 | A0AC58SGR7 | *N. tabacum* | 107780231 | GCF_000715075.1 | 982 |
| HvXRN4-like 1 | A0A8I6XLQ4 | *H. vulgare* | 123452664 | GCF_904849725.1 | 1,014 |
| ZmXRN4-like 1 | A0A1D6GFM9 | *Z. mays* | 100384297 | GCF_902167145.1 | 990 |
| AtXRN2 | Q9FQ02 | *A. thaliana* | 834261 | GCF_000001735.4 | 1,012 |
| AtXRN3.1 | Q9FQ03-1 | *A. thaliana* | 843900 | GCF_000001735.4 | 1,020 |
| AtXRN3.2 | Q9FQ03-2 | *A. thaliana* | 843900 | GCF_000001735.4 | 763 |
| NtXRN3-like 1 | A0AC58TFP4 | *N. tabacum* | 107809493 | GCF_000715075.1 | 1,112 |
| HvXRN3-like 1 | A0A8I6X4N0 | *H. vulgare* | 123444770 | GCF_904849725.1 | 1,049 |
| ZmXRN3-like 1 | K7UB09 | *Z. mays* | 100501093 | GCF_902167145.1 | 1,058 |
| ZmXRN3-like 2 | A0A1D6MQ11 | *Z. mays* | 103649964 | GCF_902167145.1 | 1,066 |
| HvXRN-like 1 | A0A8I7BEU4 | *H. vulgare* | HORVU.MOREX.r3.5HG0492360 | GCF_904849725.1 | 810 |
| ZmXRN-like 1 | A0A804NZB2 | *Z. mays* | 103654383 | GCF_902167145.1 | 893 |
| FaXRN-like | A0A7D3QU49 | *F. algeromassiliense* | MT418680 | GCF_029885335.1 | 595 |

Table S 2. Plasmids used in this thesis.

| **Name** | **Construct** | **Usage** | **Source** |
| --- | --- | --- | --- |
| #0533 pRAK_133 | Leader-xrRNA5 Level 0 | IVT of Leader-xrRNA5 | Working group Koch |
| #594 pRAK_140 | Leader-xrRNA5 mut Level 0 | IVT of Leader-xrRNA5mut | Working group Koch |

Table S 3. Oligonucleotides used in this thesis.

| **Stock ID** | **5′–3′ sequence** | **Usage** | **Source** |
| --- | --- | --- | --- |
| #1070 T7-Leader xrRNA5 For. | GATCGGATCCTAATA  CACTCACTATAGGAA  GAATGAAACTGTCACC | IVT of Leader-xrRNA5 and  Leader-xrRNA5mut | Working group Koch |
| #1071 ST9a rev. | TGCCTACAGGACTGA  TAGCC |  |  |

^a)^ Primers were designed using SnapGene 8.2 (available at <https://www.snapgene.com/>: last accessed 21.07.2026)

Table S 4. Amino acid positions of the most interacting amino acids.

| **Enzyme** | **Glu** | **Structurally neighboring  Asp residues** | | **Asp** | **Asp  Tower domain** | **His** | **Trp** |
| --- | --- | --- | --- | --- | --- | --- | --- |
| DmXRN1 | 177 | 205 | 207 | 288 | 86 | 41 | 540 |
| ScXRN1 | 178 | 206 | 208 | 291 | 86 | 41 | 638 |
| ScXRN2 | 205 | 233 | 235 | 333 | 102 | 59 | 732 |
| AtXRN4 | 206 | 234 | 236 | 342 | 104 | 61 | 573 |
| NtXRN4-like 1 | / | 234 | 236 | / | / | 61 | 573 |
| NtXRN4-like 2 | 206 | 234 | 236 | 346 | / | 61 | 573 |
| NbXRN4-like 1 | 206 | 234 | 236 | 346 | / | 61 | 573 |
| HvXRN4-like 1 | 206 | 234 | 236 | / | 104 | 61 | 571 |
| ZmXRN4-like 1 | 206 | 234 | 236 | / | / | 61 | 571 |
| AtXRN2 | 207 | / | 237 | 346 | 105 | 62 | 659 |
| AtXRN3.1 | 205 | 233 | 235 | 346 | / | 60 | 667 |
| AtXRN3.2 | 205 | 233 | 235 | 346 | 103 | 60 | 667 |
| ZmXRN3-like 1 | 205 | 233 | / | / | 103 | 60 | 669 |
| HvXRN-like 1 | 199 | 227 | 229 | 340 | 95 | 49 | 624 |
| ZmXRN-like 1 | 210 | 238 | 240 | 346 | 106 | 62 | 641 |
